# Shedding light on blue-green photosynthesis: A wavelength-dependent mathematical model of photosynthesis in *Synechocystis* sp. PCC 6803

**DOI:** 10.1101/2023.06.30.547186

**Authors:** Tobias Pfennig, Elena Kullmann, Tomáš Zavřel, Andreas Nakielski, Oliver Ebenhöh, Jan Červený, Gábor Bernát, Anna Matuszyńska

## Abstract

Cyanobacteria hold great potential to revolutionize conventional industries and farming practices with their light-driven chemical production. To fully exploit their photosynthetic capacity and enhance product yield, it is crucial to investigate their intricate interplay with the environment including the light intensity and spectrum. Mathematical models provide valuable insights for optimizing strategies in this pursuit. In this study, we present an ordinary differential equation-based model for the cyanobacterium *Synechocystis* sp. PCC 6803 to assess its performance under various light sources, including monochromatic light. Our model can reproduce a variety of physiologically measured quantities, e.g. experimentally reported partitioning of electrons through four main pathways, O_2_ evolution, and the rate of carbon fixation for ambient and saturated CO_2_. By capturing the interactions between different components of a photosynthetic system, our model helps in understanding the underlying mechanisms driving system behavior. Our model qualitatively reproduces fluorescence emitted under various light regimes, replicating Pulse-amplitude modulation (PAM) fluorometry experiments with saturating pulses. Using our model, we test four hypothesized mechanisms of cyanobacterial state transitions. Moreover, we evaluate metabolic control for biotechnological production under diverse light colors and irradiances. By offering a comprehensive computational model of cyanobacterial photosynthesis, our work enhances the basic understanding of light-dependent cyanobacterial behavior and sets the first wavelength-dependent framework to systematically test their producing capacity for biocatalysis.

## Introduction

Cyanobacteria are responsible for a quarter of global carbon fixation [1]. They are, in fact, the originators of oxygenic photosynthesis, later transferring this capability to other organisms via endosymbiosis [2]. Despite their relative simplicity in cellular structure, the cyanobacterial photosynthetic machinery is a highly sophisticated system that shows significant differences from their plastidic relatives [3]. Recently, they have emerged as a powerful resource for research and biotechnology due to their unique combination of beneficial traits and photosynthetic capabilities [4]. In the quest for environmentally friendly alternatives to fossil fuels and sugar-based production, cyanobacteria stand out as promising candidates due to their ability to convert sunlight and CO_2_ into valuable products, their minimal growth requirements, and their adaptability to diverse environments. Their metabolic versatility allows for producing a wide range of biofuels, chemicals, and raw materials. Besides biomass [5], the cells can be harvested for a variety of primary and secondary metabolites, such as sugars and alcohols [6, 7], chlorophyll and carotenoids [4], (poly)peptides and human vitamins [8], and terpenoids [9]. In particular, strains of the model cyanobacteria *Synechocystis* sp. PCC 6803 and *Synechococcus elongatus* PCC 7942, are highly attractive platform organisms for the phototrophic production of e.g. isoprene, squalene, valencene, cycloartenol, lupeol or bisabolene [9]. Leveraging the cells’ natural capabilities, isolation of molecular hydrogen [10] and reduced nitrogen [4] is also possible, with uses in energy and agronomic sectors. Furthermore, there have been attempts to use cyanobacteria for bioelectricity production [7, 11] or, inversely, to overcome cellular limitations by fuelling cyanobacteria with induced electrical currents [12].

Modifying metabolism for biotechnological purposes involves overcoming natural regulations and inhibitory mechanisms, disrupting the metabolic network’s balance. However, balance is crucial for a proper photosynthetic function [13] and, thus, the viability of cyanobacteria for biotechnology. Therefore, a comprehensive understanding of primary and secondary metabolism is essential for effective and compatible modifications. Mathematical models integrate and condense current knowledge to identify significant parts and interactions, enabling the simulation of the effect of various external factors and internal modifications [14, 15]. They can also provide a platform to test new hypotheses. Numerous plant models of primary metabolism helped to identify the most favorable environmental conditions, nutrient compositions, and genetic modifications to maximize the desired outputs [16, 15]. Despite the evolutionary connection between cyanobacteria and plants, the structural and kinetic differences between cyanobacteria and plants (e.g., competition for electrons due to respiration [17], phycobilisomes (PBSs) as cyanobacterial light-harvesting antennae, photoprotection mediated by Orange Carotenoid Protein (OCP), existence of Carbon Concentrating Mechanism (CCM)) prevent the use of established plant-based models for photosynthesis [3, 17, 18, 19, 20, 21]. Even standard experimental methods developed for plants for non-invasive probing of photosynthesis using spectrometric techniques, such as Pulse Amplitude Modulation (PAM) fluorometry and the Saturation Pulse method (PAM-SP)[22], may require either adaptation or change in the interpretation of the measurements when applied to cyanobacteria [3, 23, 24]. In PAM fluorometry, a modulated light source is used to excite the chlorophyll molecules [22]. The emitted fluorescence is then measured, and various parameters derived from this fluorescence signal can provide insights into the efficiency of photosynthesis, the health of the photosynthetic apparatus, and other aspects of plant physiology. Compared to plants and green algae, the measured fluorescence of cyanobacteria has contributions from PSII, PSI, and detached PBS resulting in distinct fluorescence behavior [3, 24, 25, 26].

Therefore, a mathematical model targeted specifically for cyanobacteria, and capable of simulating and interpreting their re-emitted fluorescence signal after various light modulations is needed to obtain a system perspective on their photosynthetic dynamics. Established cyanobacterial models often describe broad ecosystem behavior or specific cellular characteristics [27]. Worth mentioning here are constrained-based reconstructions of primary metabolic networks [28, 29, 30], as well as kinetic models, ranging from simple models of non-photochemical quenching [31] and fluorescence [26] to adapted plant models to study the dynamics of cyanobacterial photosynthesis [32] and models created to study proteome allocation as a function of growth rate [33]. However, none of these models provide a detailed, mechanistic description of oxygenic photosynthesis in *Synechocystis* sp. PCC 6803, including the dynamics of respiration and a mechanistic description of short-term acclimation mechanisms, which are highly sensitive to changes in light wavelengths.

With this work, we provide a detailed description of photosynthetic electron flow in cyanobacteria (as summarized in Fig. 1a), parameterized to experimental data, including measurements collected under monochromatic light, in *Synechocystis* sp. PCC 6803, a unicellular freshwater cyanobacterium. Light is a critical resource for photosynthetic prokaryotes, which defines their ecological niche and heavily affects cell physiology [24, 34, 35]. Significantly, beyond its intensity, the light spectrum plays a crucial role in exerting physiological control. For example, growth under various monochromatic light sources led to large differences in cyanobacterial growth rate, cell composition, and photosynthetic parameters [36]. Blue light strongly inhibits growth and can cause cell damage by disrupting the excitation balance of photosystems [29, 37, 38], resulting from the varying absorption properties of their pigments [38]. To react to changes in illumination, the cell is able to undergo both short and long-term adaptations. Over time, cells adjust their pigment content (in a process called chromatic acclimation), and the ratio of photosystems to optimize performance [35, 24]. In the short term, processes like OCP-related Non-Photochemical Quenching (NPQ) [39, 40] or state transitions [41] help them adapt, though precise mechanisms of the latter are not yet fully elucidated [3, 42]. While the scientific community agrees that the PQ redox state triggers state transitions, multiple underlying mechanisms have been proposed without a current consensus. Our model uses therefore both light intensity and light wavelengths as input, allowing the simulation of any combination of light sources and adaptation to the specific growth conditions. Readouts include all intermediate metabolites and carriers, most importantly ATP and NADPH, fluxes through several electron pathways: Linear Electron Transport (LET), Respiratory Electron Transport (RET), Cyclic Electron Transport (CET) and Alternate Electron Transport (AEF)), reaction rates, such as carbon fixation and water splitting, and the cell’s emitted fluorescence as measured by PAM. We perform Metabolic Control Analysis (MCA) [43, 44, 45] of the network in different light conditions, showing that the reactions which determine the rate of CBB flux shift from photosynthetic source reactions to sink reactions within the CBB as light intensity increases. By harnessing the power of mathematical modelling, we seek to provide a quantitative framework to test further hypothesis on the photosynthetic mechanisms in cyanobacteria and contribute to basic research on these organisms that eventually can lead to optimized cyanobacterial production and contribute to the advancement of green biotechnology.

**Fig. 1:**
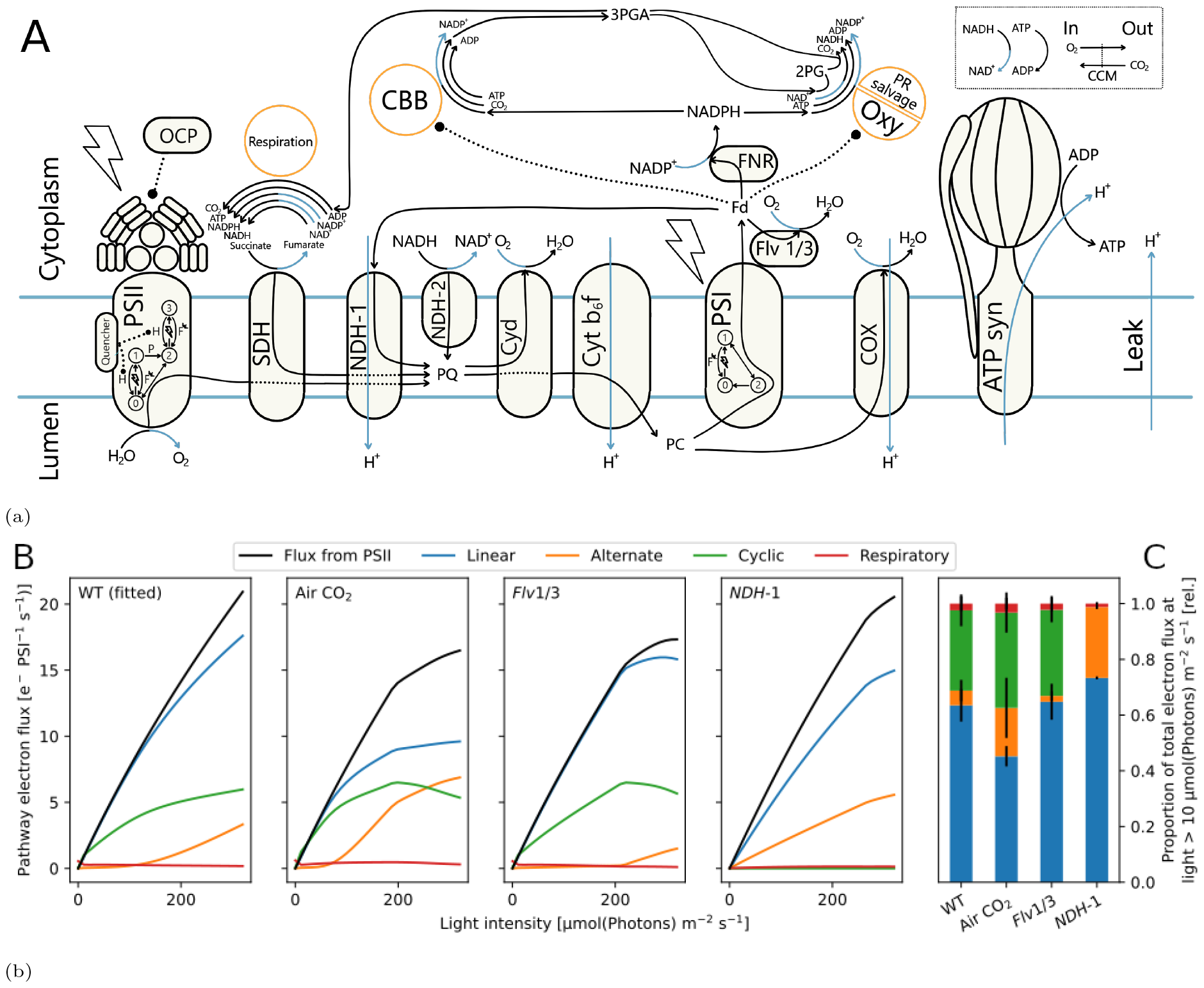
Computational model of the photosynthetic and respiratory chain allows simulating electron fluxes through main (linear), cyclic, alternate, and respiratory pathways in *Synechocystis* sp. PCC 6803. **a)** Schematic representation of components and reactions included in the model of cyanobacterial photosynthesis. The model includes descriptions of four protein complexes (PSII, PSI, Cb_6_f and ATP synthase), main electron carriers and reactions through them, enabling simulation of electron transfer through Linear Electron Transport (LET), Cyclic Electron Transport (CET), Respiratory Electron Transport (RET), and Alternate Electron Transport (AEF). With orange circles (Respiration, CBB, and photorespiratory salvage pathway (PR salvage)) we mark pathways represented in the model as lumped reactions. The top-right box shows gas exchange reactions (O2 export and active CO2 import) and metabolic ATP and NADH consumption. Electron and proton flows are colored black and blue, respectively. Regulatory effects, such as Fd-dependent CBB activity, are represented with dotted lines. The two photosystems are described using Quasi-Steady-State (QSS) approximation. For the analyzes we assume internal quencher as the state transition mechanism, as marked on PSII. Various scenarios of PBS attachment can be simulated, on the figure attached to PSII. **b)** Simulated steady-state electron flux through linear (blue), cyclic (green), alternate (flavodiiron+terminal oxidases, orange) and respiratory (red) electron pathways for light intensities between 10 µmol(photons) m^*−*2^ s^*−*1^ and 300 µmol(photons) m^*−*2^ s^*−*1^. The model has been **parameterized** to yield approximately 15 electrons PSI^*−*1^ s^*−*1^ linear electron flow (blue) for a fraction of 60 % under saturating CO2 conditions, as measured in wild type (WT) [46]. The model **predicts** flux distributions under sub-saturating air CO2 level and for the flavodiiron (*Flv* 1/3) and NAD(P)H Dehydrogenase-like complex 1 (NDH-1) knockout mutants. Each value represents a steady-state flux under continuous light exposure. Simulations were run using 670 nm light (Gaussian LED, *σ* = 10 nm). The barplot shows the mean flux distribution for light intensities over 10 µmol(photons) m^*−*2^ s^*−*1^. Error bars show sd. Abbreviations: 2PG: 2-phosphoglycolate, 3PGA: 3-phosphoglycerate, ADP: Adenosine diphosphate, ATP: Adenosine triphosphate, ATPsynth: ATP synthase, CBB: Calvin-Benson-Bassham cycle, CCM: Carbon Concentrating Mechanism, COX(aa3): Cytochrome c oxidase, Cyd(bd): Cytochrome bd quinol oxidase, Cyto b6 f: Cytochrome b6 f complex, FNR: Ferredoxin-NADP^+^ Reductase, Fd: Ferredoxin, NADP^+^: Nicotinamide adenine dinucleotide phosphate, NADPH: reduced Nicotinamide adenine dinucleotide phosphate, NAD^+^: Nicotinamide adenine dinucleotide, NADH: reduced Nicotinamide adenine dinucleotide, NDH-2: NAD(P)H Dehydrogenase complex 2, OCP: Orange Carotenoid Protein, Oxy: RuBisCO oxygenation, PC: Plastocyanine, PQ: Plastoquinone, PR: Photorespiration, PSI: Photosystem I, PSII: Photosystem II, SDH: Succinate dehydrogenase

## Methods

### Model description

We developed a dynamic, mathematical model of photosynthetic electron transport in *Synechocystis* sp. PCC 6803 (further *Synechocystis*) following a classical bottom-up development cycle. Our model consists of a system of 17 coupled Ordinary Differential Equations (ODEs), 27 reaction rates, and 95 parameters, including measured midpoint potentials, compound concentrations, absorption spectra, and physical constants (Table S3). By integrating the system over time, we can simulate the dynamic behavior of rates and concentrations of all reactions and reactants visualized on Fig. 1a and summarized in Table S2, including dynamic changes in the lumenal and cytoplasmic pH. We included a detailed description of four commonly distinguished electron transport pathways: LET, CET, RET, and AEF. Given the high similarity between the essential electron transport chain proteins of plants and cyanobacteria [3, 20], the photosystems were described using Quasi-Steady-State (QSS) approximation, as derived in our previous dynamic models of photosynthetic organisms [47, 48]. We followed a reductionist approach simplifying many downstream processes into lumped reactions. The lumped CBB, PR, and metabolic consumption reactions represented the main cellular energy sinks. Functions describing CBB and RuBisCO oxygenation (Oxy) contained multiple regulatory terms, including gas and metabolite concentrations (see SI). Although cyanobacteria CCM components include at least four modes of active inorganic carbon uptake [49], we have decided to represent the mechanism as a one lump reaction. By calculating the dissolved CO_2_ concentration at the cellular pH and with an actively 1000-fold increased intracellular CO_2_ gas pressure (see Supplement S1.12) we reflect the very efficient cyanobacterial concentrating mechanism. Unless stated otherwise, simulations were run under three assumptions: 25 *?*C temperature with 230 µmol L^*−*1^ dissolved O_2_ and supplemented with 5 % CO_2_. The pigment content and photosystems ratio were parametrized to a cell grown under ambient air with 25 µmol(photons) m^*−*2^ s^*−*1^ of 633 nm light. All rates and concentrations have been normalized to the chlorophyll content (4 mmol L^*−*1^).

The default initial metabolite concentrations were set to literature measurements (Table S1). Steady-state simulations were run for 1 *×* 10^6^ s. For the steady-state simulations, we considered that the steady-state is reached if the Euclidean norm of relative concentration changes between the last two time steps did not exceed 1 *×* 10^*−*6^. Because the regulatory processes, CBB redox activation, OCP activation, and state transitions, have slow rates of change, we compared their relative changes to a threshold of 1*×*10^*−*8^.

### Code implementation

The model has been developed in `Python` [50] using the modeling package `modelbase` [51] further exploring a highly modular approach to programming mathematical models. The model and scripts used to numerically integrate the system and to produce all figures from this manuscript, as well as analysis run during the peer review process are accessible at https://github.com/Computational-Biology-Aachen/synechocystis-photosynthesis-2024.

### Model Parameterization

The model has been manually parameterized, integrating physiological data and dynamic observations from numerous groups (pH-ranges: [52], NADPH reduction: [53], O_2_ change rates: [25], CO_2_ consumption: [33], PQ reduction: [54], PC, PSI, and Fd redox-states: [55], PAM-behavior: [42], electron fluxes: [46], PAM fluorescence: [56]).

The model depends on 96 parameters (Table S3). 43 parameters, including pigment absorption spectra, were taken directly from the literature, six parameters describe the experimental setup (light: intensity and spectrum, CO_2_, O_2_, concentration of cells, and temperature), and eight parameters describing photosystems concentrations and pigment composition were estimated from provided data [56]. The latter parameters were measured spectrophotometrically and through 77 K fluorescence, assuming a 10-times higher fluorescence yield of free PBS (compare [26]). PAM-SP fluorescence curves were used to fit seven fluorescence-related parameters, including quenching and OCP constants [56], and two parameters were fit to electron transport rate measurements [46]. Nine rate parameters were estimated from reported rates of the reaction itself or connected processes such as O_2_ generation for respiration. To derive rate constants, we divided the determined rate by the assumed cellular substrate concentrations. Five parameters stemmed from simplifying assumptions regarding inhibition constants, the cytoplasmic salinity, and pH buffering. 16 further parameters were fitted to reproduce literature behavior such as cellular redox states or regulation of the CBB.

To avoid overfitting the parameters to a particular experimental set-up, we avoided using sophisticated fitting algorithms and instead proceeded with manual curation. At every step of model refinement we have been comparing visually how the change affects our simulated redox state, oxygen evolution, carbon fixation, and dynamics of implemented photoprotective mechanisms. A comprehensive list of all the model parameters utilized in this study, including values needed for unit conversion, is provided in Table S3 and Table S4 (state transition analysis separate, see below), ensuring transparency and reproducibility of our computational approach.

### Reaction kinetics

Following the principle of parsimony, all reactions where no additional regulatory mechanism was known have been implemented with first-order Mass-Action (MA) kinetics. A reaction with substrates S_i_ and products P_j_ is defined as: ∑ *n*_*i*_*S*_*i*_ *↔* ∑ *m*_*j*_*P*_*j*_ with *i, j* ∈ ℕ where *n*_*i*_ and *m*_*j*_ are the stoichiometric coefficients of substrates and products, respectively. For each reaction, we calculated the Gibbs free energy (Δ_r_G′, see supplemental information) [47, 48]. Only reactions with Δ_r_G′ close to 0 under physiological conditions were described with reversible kinetics. Thus, we set reactions as irreversible except for ATP synthase, SDH, FNR, regulatory variables (e.g. CBB activation), and PSII and PSI internal processes.

To simplify higher order reversible MA equations [57], we first decompose the rate equation into separate kinetic and thermodynamic components (as done for Michaelis-Menten (MM) [58]) and then simplify only the kinetic part, leading to (see Equation S3 and S5):

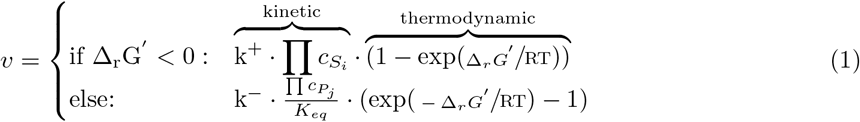

with substrate concentrations c_S_, product concentrations c, and *K*_*eq*_ = exp(*−*Δ_*r*_*G’*^*0*^ */*RT). Here, we approximate 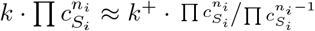 and 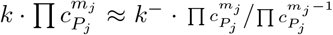 which, for any *n*_*i*_ *>* 1 or *m*_*j*_ *>* 1, leads to k^+^ ≠ k^*−*^ (see Equation S6). This necessitated parameterising k^+^ and k^*−*^ separately. The reactions FNR and SDH, which were deemed reversible, used rate Equation 1 with the determination of Δ_*r*_*G*′ during simulation. We calculated electron pathway fluxes from the following involved reactions: LET (FNR), CET (NDH-1), respiration (SDH, NDH-2), and AEF (Cyd, COX, Flv) (see Equation S47).

### Implementation of monochromatic and polychromatic light sources

To consider the influence of light spectrum on photosynthetic activity, our model takes light as input (*I*) with wavelengths (*λ*) in range between 400 and 700 nm. In this work we performed simulations using the solar spectrum, a fluorescent lamp, cold white LED, warm white LED, and “gaussian LEDs” simulated as Gauss curves with a set peak-wavelength and variance of 10 nm or 0.001 nm (“near monochromatic”) [59] (see Fig. 3a & Fig. S11).

For calculation of absorbed light we further differentiate between the light absorbed by PSI, PSII, and PBS, based on their reported pigment composition [59] (Fig. 3a). We focused on four most abundant pigments: chlorophyll, beta-carotene, phycocyanin, and allophycocyanin, although the implementation allows for more complex composition. We assume that PBSs can be either free, in which case the excitation is lost, or attached to one of the photosystems to transfer their excitation energy. The respective fractions of PBS states were fixed to values from [56], except for simulations of state transition mechanisms which required dynamic PBS behavior. We assumed pigment content and PBS-attachment as measured by Zavřel *et al*. [56], although different pigment composition can be provided as an input to the model. We calculate PSII excitation rate *E*_*PSII*_ (and *E*_*PSI*_ analogically) as:

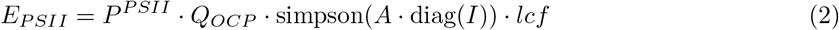

where vector *P*_1*×*4_ denotes absorbed photons redistributed to either photosystem, accounting e.g. for their pigment composition, high PSI:PSII ratio and the PBS attachment; *Q*_*OCP*_ = diag(1, 1, 1*−OCP*, 1*−OCP*) is a diagonal matrix with values set to one everywhere but at the contribution of PBS to reduce the excitation rate by light energy quenched due to OCP activity; *A*_*p×λ*_ = (*a*_*λ,p*_) contains each pigment *p*’s-specific absorption spectra; “simpson” is a row-wise, numerical integral of the light absorbed by each considered pigment, calculated using the composite Simpson’s rule (we used `scipy.integrate.simpson` function); *lcf* = 0.5 is the light conversion factor to estimate the amount of generated excitations, which was fitted to match the electron transfer rates of Theune *et al*. [46] (Fig. 1b). Importantly, we assume that despite the wavelength-dependent energy content, all photons result in equivalent excitation of photosystems with the extra energy being lost as heat [60, 61]. This implementation enabled us to simulate various light-adapted cells by updating the parameters corresponding to measured pigment composition and photosystem ratio. For simulations of PAM-SP, we further calculate the light encountered by a mean cell (*I*) for each wavelength according to an integrated Lambert-Beer function [62] accounting for the decreasing irradiance at various depths due to cellular absorption (see SI Equation S67).

### Activation of photosystems

Following our previous approach [47], we modeled the photosystem’s excitation and internal electron transport assuming that i) they are at Quasi-Steady-State (QSS), as they operate on a much faster time scale than other photosynthetic reactions, and ii) at every time point, photosystem II can be in one of four possible states, and PSI in three, relating pigment excitation with the charge separations at reaction centers (Fig. S1). The PSII excitation rate constant *k*_*LII*_ is calculated from *E*_*PSI*_ in Equation 2 (in µmol(photons) mg(chl)^*−*1^) multiplied by the molar mass of chlorophyll *M*_*Chl*_ and normalized to PSII concentration:

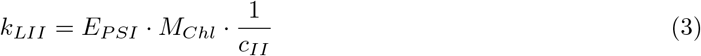

PSII was described with four (*B*_0_ - *B*_3_) and PSI with three states (*Y*_0_ - *Y*_2_) (see Fig. 1a and Fig. S1). The QSS models also consider the relaxation of excitations by fluorescence or heat emission (only PSII). We defined the PSII rate *v*_*PSII*_ as

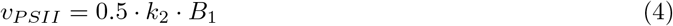

since two *B*_1_ *→ B*_2_ reactions have to occur for a full PQ reduction.

### Calculating the fluorescence signal

Based on the principle of PAM measurements, the model calculates fluorescence proportional to the gain in excited internal states of PSII and PSI when adding measuring light to the growth light. Additionally, we consider fluorescence of free PBS using their light absorption [26]. The default measuring light is set to 625 nm at 1 µmol(photons) m^*−*2^ s^*−*1^ throughout this manuscript.

PAM fluorometry measures the cellular fluorescence emitted in response to microsecond pulses of measuring light with a constant, low intensity. We built our model fluorescence function on the same principle. Measuring light pulses with irradiance *I*_*ML*_ are applied on top of the actinic light *I*, so cells experience a total irradiance of *I*_+*ML*_ = *I* + *I*_*ML*_. We then recalculate the photosystems’ QSS systems using *I*_+*ML*_, resulting in the internal states 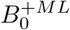 to 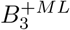 and 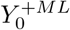 to 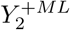 We then define the PAM fluorescence as the increase in photosystem fluorescence reactions by the addition of measuring light (see Fig. S1):

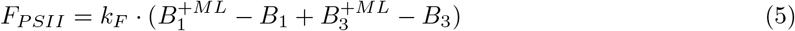

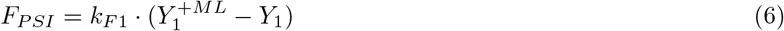

Lastly, we make the simplifying assumption that PBS fluorescence only results from the fraction of uncoupled PBS 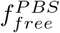 and is proportional to their absorption of *I*_+*ML*_:

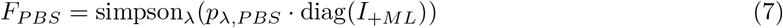

Considering cyanobacterial optical properties of light-harvesting pigments, fluorescence measured at room temperature can originate from both photosystems and PBS [3]. There have been attempts to determine the fluorescence contributions of each component [26]. We assume that the three fluorescent species contribute differently to the fluorescence detected *>* 650 nm, e.g. because of differing emission spectra. Therefore, we include weighing factors when calculating the total recorded fluorescence in Equation 8, which were calculated in Fig. 2a fitting. We estimate the total measured fluorescence signal by calculating the weighted sum of PSI, PSII, and PBS fluorescence:

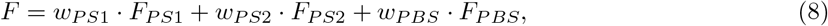

where weights *w* were manually fitted to reproduce the fluorescence dynamics under changing light conditions of the experiment displayed in Figure 3 (fitted values can be found in (Table S3) as `fluo influence`).

**Fig. 2:**
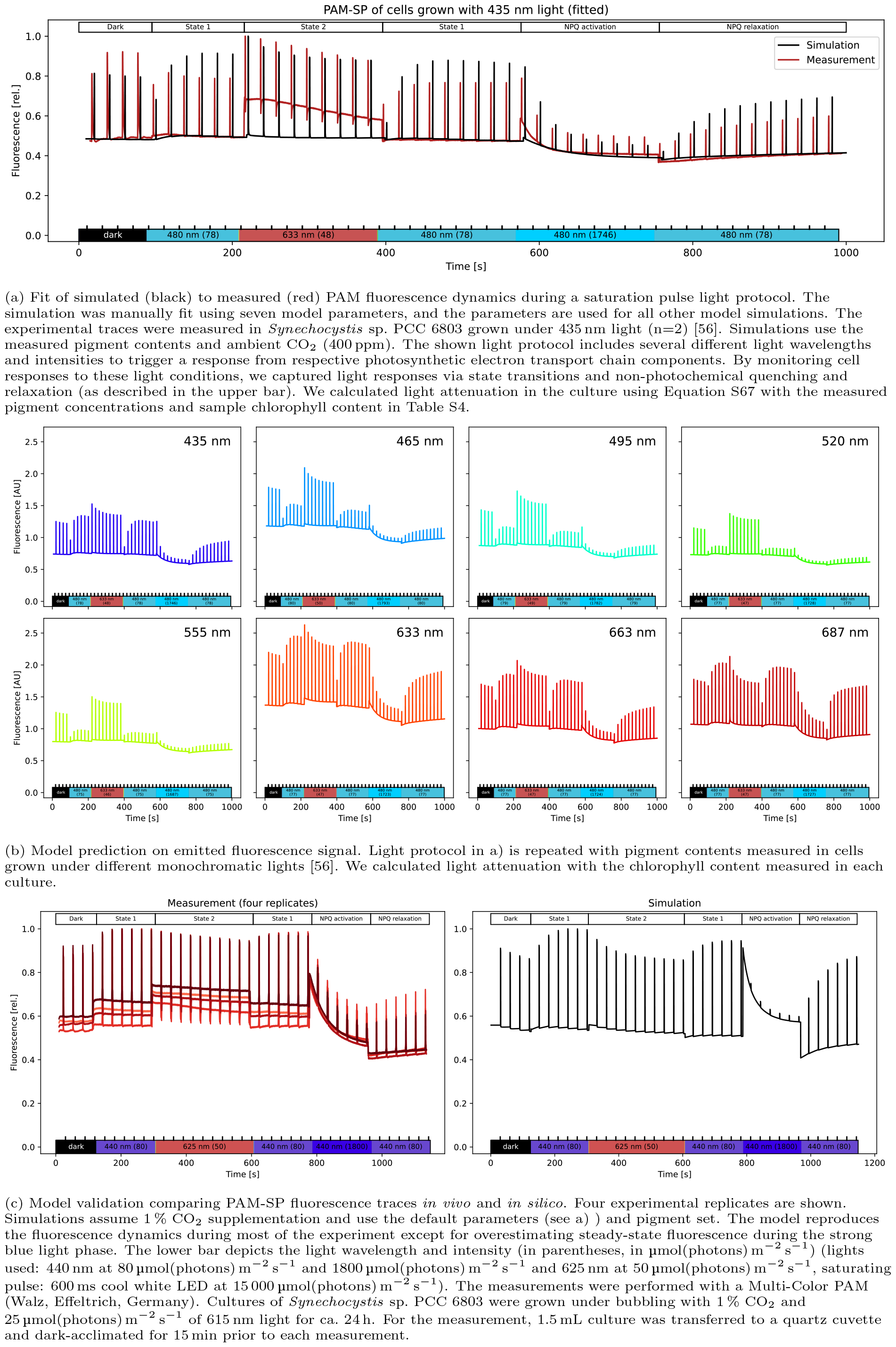
Saturation pulse method using Pulse Amplitude Modulation (PAM) fluorescence measurement *in vivo* and *in silico*. The simulated signal has been calculated using (Equation 8). All experimental measurements were performed with a Multi-Color PAM (Walz, Effeltrich, Germany).

**Fig. 3:**
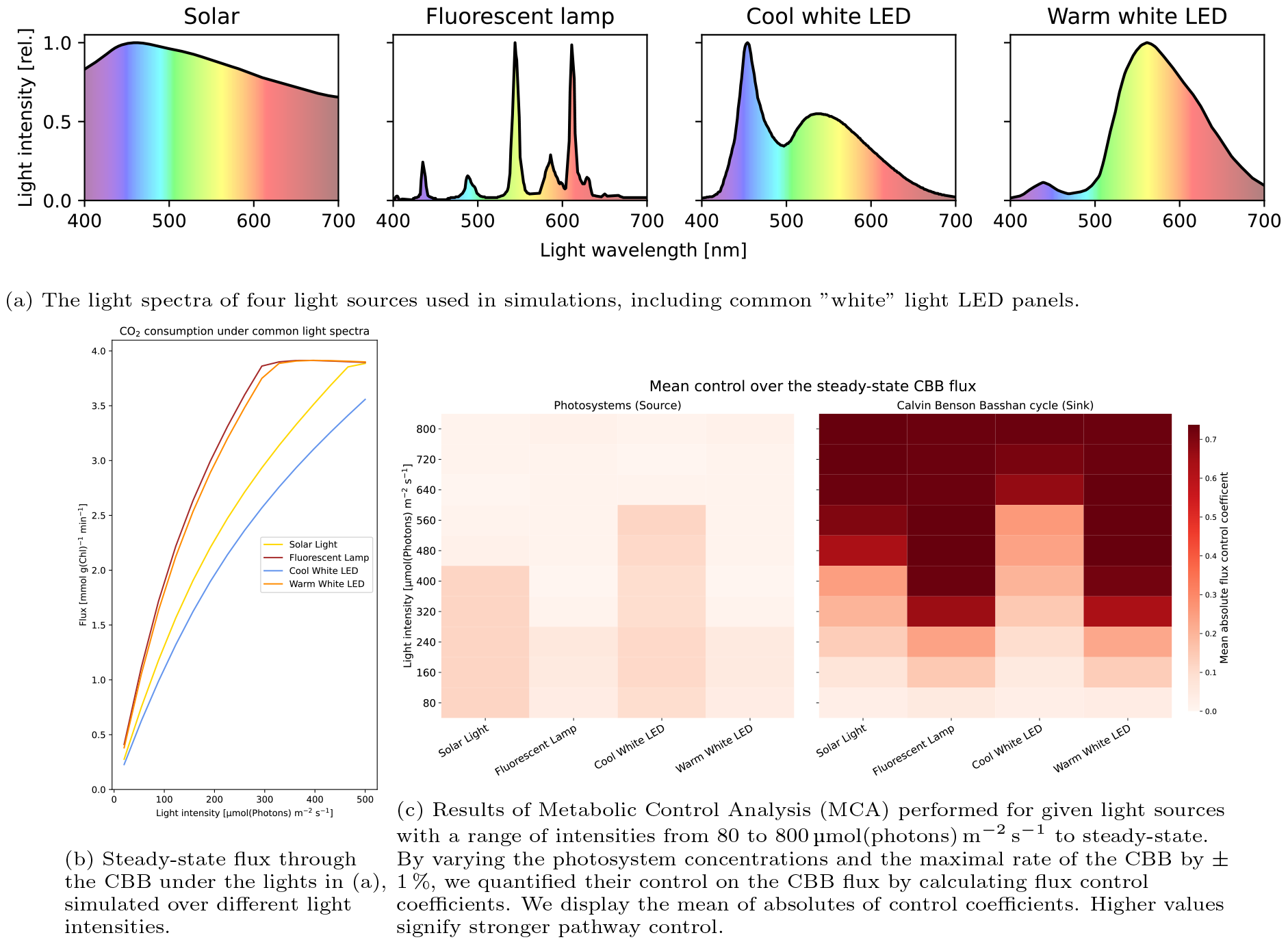
Systematic analysis of the effect of various light sources on the rate of carbon fixation.

### Testing four possible mechanisms of state transitions

We intended to use the model to provide quantitative arguments for a possible mechanism of state transitions that is not yet fully elucidated. We have implemented and tested four proposed state transition mechanisms based on a recent review [42] (Fig. 4a.) We model the transition to state 2 depending on reduced PQ (*PQ*_*red*_) and to state 1 on oxidized PQ (*PQ*_*ox*_). We implemented the default PSII-quenching (used for simulations in Fig. 2a) using a constitutively active quenching reaction and a reverse reaction with Hill kinetics. The remaining state transition models were described with few reactions and using MA kinetics. A complete mathematical description of the implementations is available in the SI. For the analysis, we systematically varied the parameter sets of all implementations and compared the steady-state fluorescence and PQ redox state under different lighting conditions (actinic: 440 nm at 80 µmol(photons) m^*−*2^ s^*−*1^ or 633 nm at 50 µmol(photons) m^*−*2^ s^*−*1^; measuring: 625 nm at 1 µmol(photons) m^*−*2^ s^*−*1^) (parameters in Table S5).

**Fig. 4:**
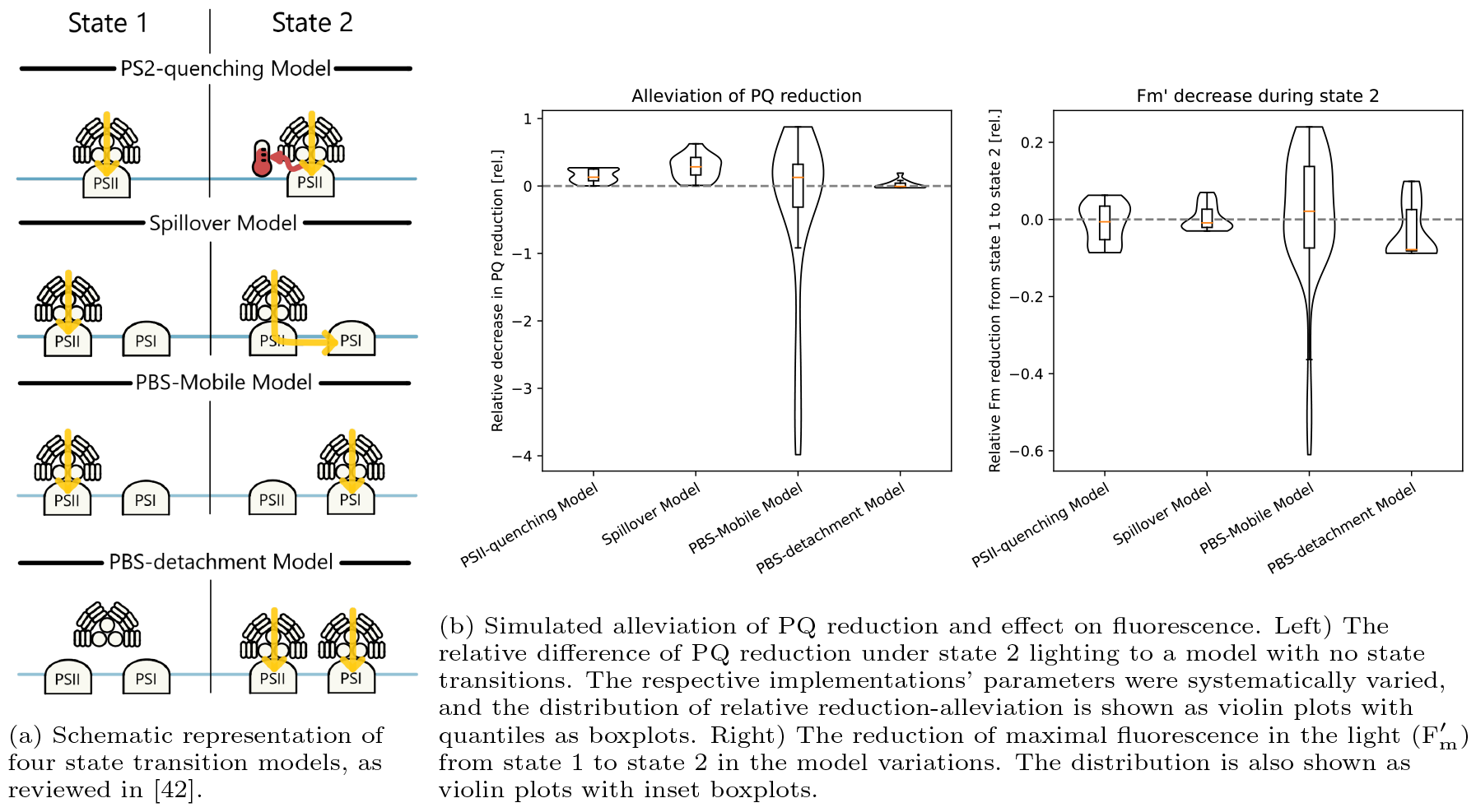
Testing possible mechanisms of state transitions. Four possible mechanisms of state transitions have been implemented and parameterized randomly to quantify their performance by contributing to oxidising PQ pool. The fluorescence signal has been calculated using (Equation 8).

### Metabolic Control Analysis

Metabolic Control Analysis (MCA) is a quantitative framework to study how the control of metabolic pathways is distributed among individual enzymes or steps within those pathways. It quantifies the change in steady-state compound concentrations or reaction fluxes in response to perturbation of an examined reaction [43, 45]. We used the `modelbase.mca` function `get response coefficients df` to perform MCA on our model. The function is using definitions proposed by [63, 43] and calculates the flux control coefficients 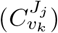 and concentration control coefficients 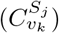 using formulas:

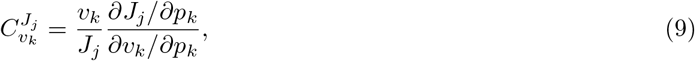

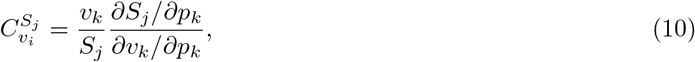

where *J*_*j*_ and *S*_*j*_ are respectively the steady-state fluxes and concentrations of the system, *p*_*k*_ is a kinetic parameter which affects directly only reaction *k* with the rate *v*_*k*_ (see [63, 43]). We approximated these formulas numerically using the central difference, varying the parameters by *±*1%. MCA has been repeated for various simulated irradiances (Fig. 3a). For systematic analysis of the effect of various light sources on the rate of carbon fixation, we calculated the absolute of the control coefficients and show the mean of the following sets of model reactions: Photosystems (PSI, PSII), light-driven (PSI, PSII, Cytochrome b_6_f complex, NDH-1, FNR), alternate (Flv, Cytochrome bd quinol oxidase (Cyd), Cytochrome c oxidase), and respiration (lumped respiration, Succinate Dehydrogenase, NDH-2).

### Analysis of the production capacity

Exploring highly modular structure of the model, for determining the production potential of a biotechnological compound, we simply added an irreversible model reaction consuming ATP, NADPH, and Fd in the required ratio. We assume optimality of carbon provision by the CBB and, thus, set its rate to zero and add the energy equivalent cost of carbon fixation to the cost of the biotechnological compound. The sink reaction was described using simplified MA kinetics with a rate constant sufficient to prevent substrate accumulation under any light intensity (here set to 5000 (unit depending on the order of the reaction) for every sink). Additionally, we added MA reactions draining ATP and NADPH with a very high rate constant (10 000 s^*−*1^) if their pools became filled over 95 % to avoid sink limitation by either compound.

## Results

We present the first kinetic model of photosynthesis developed for cyanobacteria that can simulate its dynamics for various light intensities and spectra. It is developed based on well-understood principles from physics, chemistry, and physiology, and is used as a framework for systematic analysis of the impact of light on photosynthetic dynamics. Our analysis focuses on several key aspects: the redox state of electron carriers, carbon fixation rates in ambient air, reproduction of fluorescence dynamics under changing light conditions, and the electron flow through main pathways (LET, CET, AEF) under different conditions. We used O_2_ measurements [64] to qualitatively validate the PSII light harvesting and photochemistry implementation of the model. Changes in our simulated steady-state O_2_ evolution rates are in quantitative agreement with the experimental data, during low light and exceed measured rates under light saturation by ca. 20 % (Fig. S4). Unfortunately, the exact culture conditions (e.g. density) and strain used in the reference work [64] are not known. Hence the pigment composition may differ. We also calculated the fraction of open PSII for increasing light intensities to assess the model quality (Fig. S5). We observed that our response curve is less sensitive to increasing light, as our PSII are open for higher light intensities than reported 300 µmol(photons) m^*−*2^ s^*−*1^ [65].

### Flux through alternative electron pathways

We simulated the steady-state flux of electrons through the PETC for four transport pathways under 670 nm monochromatic illumination (Fig. 1b). We parameterized the flux through the LET to yield approximately 15 electrons PSI^*−*1^ s^*−*1^ and 60 % of the total PSI electron flux in the wild type (WT) [46]. Our simulated saturation of CET around 300 µmol(photons) m^*−*2^ s^*−*1^ compares well to proton flux measurements by Miller *et al*. [65]. Under ambient CO_2_ (400 ppm), our model simulates an overall limitation of electron flux and an increase in alternative flows. We found similar electron partitioning between WT and in the *Flv1/3* mutant at lower light intensities agreeing with the findings of Theune *et al*. [46]. However, our simulations show significant AEF in the WT over 200 µmol(photons) m^*−*2^ s^*−*1^, which might have been suppressed by high CO_2_ and pH in the experiments by Theune *et al*. (personal correspondence, see also [66]).

Under intermediate light intensity, the *Flv1/3* mutant also showed a higher CET while maintaining LET similar to the WT, pointing towards a balancing act of NDH-1. Inversely, our simulated NDH-1 mutant maintained high AEF but, in contrast to Theune *et al*., significant flux through the LET. In addition to simulating electron flow, our model can probe the intracellular redox state, pH, and additional fluxes through key biochemical reactions (Fig. S8). For example, it can be seen that a reduced PQ pool under high light leads to reduced CET mediated by NDH-1 and, in turn, a decreased CBB flux due to insufficient provision of ATP. Furthermore, we find that mutations affecting the electron flow lead to an increased Non-Photochemical Quenching (NPQ) at higher light intensities and the decrease in photosynthetic yield (Fig. S7).

### Photosynthesis dynamics captured via fluorescence measurements

Using experimental measurements (pigment concentrations, photosystem ratios, and expected PBS-attachment - method currently under review)[56], we manually fitted model parameters to represent a *Synechocystis* strain grown under 435 nm monochromatic light. With this model, we simulate fluorescence in a PAM-SP light protocol [22], which investigates photosynthesis behavior using the dark-adapted minimal (F_0_) and maximal fluorescence (F_m_), the maximal fluorescence in the light 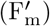 and the constantly measured steady-state fluorescence (F). By monitoring cell responses to changing light conditions, we captured light responses via state transitions and non-photochemical quenching and relaxation (for a review of the mechanisms, see [3] and for related models in plants [15]). We simulated the same light protocol of blue and red light, as used *in vivo* [56], and fit parameters controlling the fluorescence composition, state transitions, and NPQ to cells grown under 435 nm light (Fig. 2a).

Our simulation qualitatively reproduces the transition between states 1 and 2 and the activation and relaxation of NPQ by the Orange Carotenoid Protein (OCP). Because our model underestimates PSII closure in response to light, the steady-state fluorescence during light phases is also underestimated. By systematically comparing our simulation results and experimental data, we have revealed that the experimentally used saturation pulses were non-saturating in 480 nm actinic light and induced fluorescence quenching, as confirmed by follow-up experiments (see Fig. S14 in [56] and Fig. S6 in this work). Thus, we found the model’s usefulness in investigating fluorescence measurements. Using the same fitted parameters, we can also reproduce the quantitative behavior of cells grown under 633 nm monochromatic light (Fig. S3) and predict the fluorescence under further adapted pigment contents (Fig. 2b). The model shows a strong effect on cellular reactions and fluorescence when adapted to pigment contents of cells grown under other monochromatic lights. We have further validated our model against the newly measured fluorescence trace Fig. 2c. Our simulations predict accurately 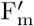, but overestimate fluorescence signal in high 440 nm light (Fig. 2c).

### Common light sources affect the metabolic control differently

Photosynthesis experiments can be conducted with many different light sources that are equivalent in photon output but differ in the spectrum. To further investigate how these spectral differences affect cellular metabolism, we simulated the model with different monochromatic and “white” light sources: solar irradiance, fluorescent lamp, and cool and warm white LED (Fig. 3a). For each light, we simulated the model to steady-state to perform MCA (Fig. 3c). We perturbed single parameters of the PETC components by ± 1 % and quantified the effect on the steady-state fluxes and concentrations. A high control coefficient represents a strong dependency of the pathway flux on changes to that parameter, with control in a metabolic network being distributed across multiple reactions. A single parameter being in full control of the flux through a network would represent the case of a typical bottleneck, but this rarely occurs in biological systems [43, 45]. We show that the electron pathway-specific control differs between the simulated light sources. Our results indicate that, at lower intensities of solar and cool white LED light, the control mainly lies within the photosystems as sources of energy carriers (Fig. 3c). We find less control by the photosystems for light spectra with a higher proportion of red wavelengths, suggesting such light sources induce less source limitation. Accordingly, the maximal simulated CO_2_ consumption is reached at lower light intensities for these spectra (Fig. 3b). All tested spectra show the CBB having the main control of CO_2_ fixation only under increased light, marking a shift towards the energy carrier sink limitation.

Repeating the analysis with simulated monochromatic lights, we found similar differences that seem to correspond with the preferential absorption by either chlorophyll or PBS (Fig. S11). The earliest switch to sink limitation was found in 624 nm light, while light that is weakly absorbed by photosynthetic pigments, such as 480 nm, seems to have little control effect. Our analysis also confirmed the intuitive understanding that remaining respiration under low light could have low control on the CBB while alternate electron flow became influential under light saturation (Fig. S10). Using the model, the control of single components, such as photosystems, can also be investigated (Fig. S12).

### Model as a platform to test alternative mechanisms of state transition

We defined four functions representing the mechanism of state transitions with the PSII-quenching model [67] as the default. Therein, a higher fraction of PSII excitations is lost as heat in state 2. Three alternative models of state transition were tested: the Spillover Model, where state 2 induces PSII excitation energy transfer to PSI; the PBS-Mobile model, where PBS attach preferentially to PSII or PSI in states 1 and 2, respectively; and the PBS-detachment Model, where PBS detach from the photosystems in state 1 [67, 42]. We model the transition to state 2 under a reduced PQ pool while oxidized PQ promotes state 1. To test the general behavior of these mechanisms without limitation to a single parameter set, we systematically simulated each mechanism with a range of parameter values, typically varying each parameter within two orders of magnitude. We calculated the relative difference of PQ reduction under state 2 lighting to a model with no state transitions, and the reduction of 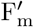 from state 1 to state 2 (Fig. 4b left). We see that all models can alleviate the PQ reduction that cyanobacteria encounter in state 2. However, across all parameter variations, the PBS-detachment model has vastly lower simulated potential to alleviate the reduction. Variations of the PBS-mobile model simulated a wide range of effects on the redox state, from near total oxidation to increased reduction. During the transition to state 2, the height of F_m_ is expected to decrease, which all models can simulate within their parameter variations (Fig. 4b right). Again, the simulations of the PBS-mobile model had the widest wide range of simulation results.

### Model as a platform to test optimal light for biotechnological exploration

Cyanobacteria show potential as cell factories for the production of terpenoids from CO_2_ or as whole-cell biocatalysts, which require different ratios of NADPH, ATP, and carbon. Several studies revealed that light availability is one of the main limitations of light-driven whole-cell redox biocatalysis [68]. With our model, we systematically analyzed the *Synechocystis* productivity for various light sources.

To identify potentially optimal light conditions and/or quantify the maximal production capacities for these exemplary processes, sink reactions were added to the model, and production was simulated with different light conditions (Fig. 5). These sinks drained the required amounts of ATP,NADPH, and Fd necessary for fixing the required amount of CO_2_ and producing one unit of the target compound. Additionally, it was necessary to add reactions that avoid overaccumulation of ATP and NADPH in case the sink was not sufficiently consuming both. The model simulates that NADPH production was highest under red (624 nm) illumination saturating around 800 µmol(photons) m^*−*2^ s^*−*1^. We also compared the simulated productions of isoprene and sucrose, which require different optimal rations of ATP and NADPH, 1.46 and 1.58, respectively. Isoprene production showed a stronger dependency on red-wavelength light, exceeding the production in blue light twofold, and did not saturate within the simulated light intensity range. Presumably, the involvement of Fd as a substrate further favors the usage of light preferentially exciting PSII. In line with the recent work by Rodrigues *et al*. (2023) [29], our simulated isoprene also follows the measured cellular growth rate as predicted by their stoichiometric model analysis (Fig. S13). Accordingly, Rodrigues *et al*. discuss that their experimentally realized isoprene production was determined not just by differential excitation of photosystems. On the other hand, simulation of the more ATP-intensive sucrose production was saturated at much lower light intensities and even decreased slightly under high light. These simulations indicate that the optimal light intensity could be lower for synthesis reactions requiring more ATP. It has also been suggested that the ATP:NADPH ratio is increased under blue light due to higher CET activity [69, 70]. However, our model did not show a benefit of more chlorophyll-absorbed light on the reactions involving ATP. Overall we found 624 nm light, to have the highest simulated production across the tested compounds and lights.

**Fig. 5:**
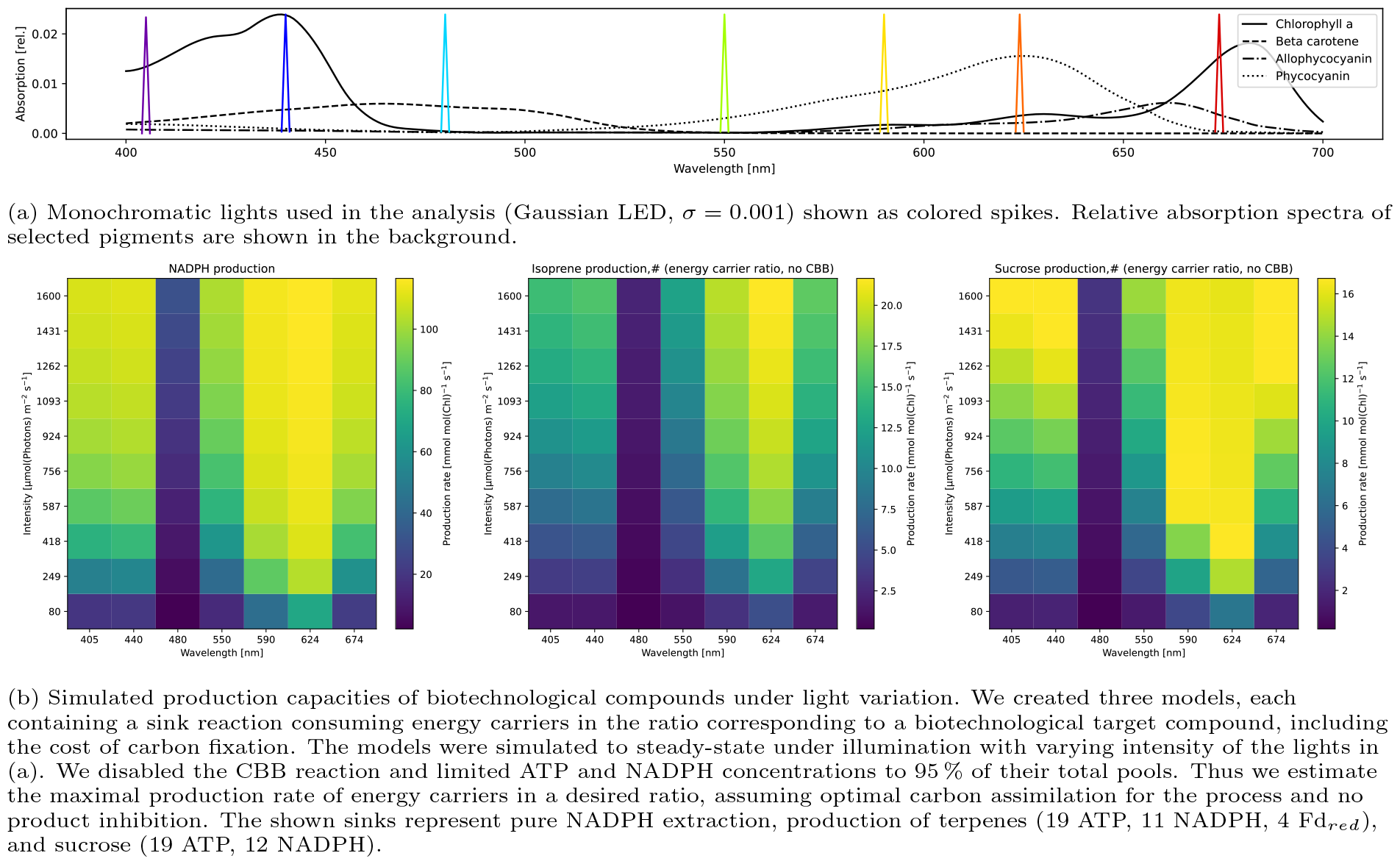
Simulated steady-state production rates of target compounds under monochromatic lights with varying intensities.

While these results were obtained for an unadapted cell, our model allows us to repeat such analyses with any adapted pigment composition (comparison of estimated CO_2_ consumptions in S9).

## Discussion

In this work, we present the first wavelength-dependent mathematical ODE-based photosynthesis model for cyanobacteria. The model contains all major processes involved in the *Synechocystis* photosynthetic electron flow, from light capture to CO_2_ fixation [17] and a description of the respiratory chain embedded within the same membrane. Furthermore, cyanobacteria-specific mechanisms were implemented in the model, including state transitions and OCP-mediated NPQ [3, 39, 71, 42]. In contrast to other existing dynamic models of photosynthesis, our model takes pigment composition of the strain as an input and can simulate illumination within the full visible spectrum (400-700 nm). Hence, results obtained with our model provide insights into the intricate dynamics of the photosynthetic process under various light conditions.

The model was validated against published measurements of gas exchange rates (Fig. S4) and compared to *in vivo* electron pathway fluxes and cellular fluorescence. The quantitative agreement with oxygen production rates supports our pigment-specific implementation of light absorption, which allows for a better assessment of the possible effect of photosystem imbalance [38, 59]. After parameterizing the model to reproduce the electron fluxes in the wild type, we used it to gain in-depth information on the system’s behavior using *in-silico* mutants. Simulations of a Flv knockout mutant showed increased CET by NDH-1 under intermediate light (Fig. 1b). It was reported previously that the proteins provide redundancy for alleviating redox stress [55, 72]. Furthermore, in the Flv mutant, flux from PSII is decreased due to lack of electron outflow to Flv (see Fig. S7). The decreased PSII flux is accompanied by raised NPQ under high light intensities.

Our calculated PAM fluorescence signal is composed of signals originating from both photosystems and PBS, with a similar contribution as in the previously published model [26]. We employed this fluorescence estimate to fit a PAM-SP experiment inducing state transitions and OCP quenching (Fig. 2a). We reached a qualitative agreement in the fluorescence dynamics, especially during the induction of OCP. Therefore, despite existence of more detailed models of OCP dynamics [31], we decided to keep our two-state implementation. The description of state transitions is challenging, as there currently is no literature consensus on the mechanism of state transitions [3, 42]. Therefore, we used our model to compare the implementations of four proposed mechanisms based on the cellular redox state and fluorescence. Simulations of all mechanisms could reproduce the expected cellular effects in some form. We saw, however, that the movement of PBS had the highest dynamic range of reducing or oxidizing the PQ pool, while PBS detachment in state 2 had a very modest effect. Therefore, the targeted movement of PBS could provide the cell with high control over its electron transport. The significance of PBS movement has been debated, however [73, 74, 75, 76], as has the spillover of energy between the photosystems [73, 74, 75]. It is noteworthy that considering solely the effect on PQ redox state, the implemented PSII-quenching model favored by Calzadilla *et al*. [42] does not have a significantly greater effect on the oxidation of PQ in our simulations. This limited PQ oxidation is in line with a model of plant photoinhibition where PSII quenching decreased PSII closure by ca. 10 % [77]. Overall, the mechanism of state transitions and its impact on photosynthetic balance remains to be evaluated.

Therefore, we used MCA to systematically study the effect of light (intensity and color) and determined the systems control on carbon fixation considering varying illumination: solar illumination, a fluorescent lamp, and cold and warm white LEDs (Fig. 3) of different intensities. The photosystems mainly controlled carbon fixation in simulations of low light intensity, which is in line with the limitation of light uptake and ATP and NADPH production as found in analyses of plant models [48]. Spectra with a high content of blue wavelength photons, which have been linked with an imbalanced excitation of PSI and PSII [37], showed a further increase in photosystems control. Indeed, blue light was found to increase PSII expression [78, 69], a cellular adaption possibly using this control. At higher light intensities, the maximum rate of carbon fixation became the main controlling factor. Thus, the strategy promising better productivity would involve increasing carbon fixation by e.g. additionally increasing the CO_2_ concentration around RuBisCO [79], engineering RuBisCO itself [80] or introducing additional electron acceptors and carbon sinks such as sucrose, lactate, terpenoids or 2,3-butanediol [81, 82, 83, 84, 85].

With the implementation of the spectral resolution, our model could also simulate cellular behavior in high cell densities (e.g. bioreactors), where the light conditions might differ throughout the culture [86]. We show that lighting in the orange-red spectrum requires the lowest intensity to saturate the photosystems, with a warm-white LED showing the same efficiency as a fluorescent light bulb, an important consideration when calculating process costs (Fig. 3b).

To showcase the biotechnological usability of this work, we analyzed the *Synechocystis* productivity for various light sources (Fig. 5). Many experimental studies have investigated optimal light colors for the production of biomass or a target compound, with most studies agreeing that white or red light is optimal for cell growth but varying results for target compounds [87, 88, 29]. Especially the synthesis of light harvesting or protection pigments is regulated and strongly dependent on the light color [89, 90, 91, 92]. These works point out that biotechnological production can be strongly improved using “correct” lighting. However, finding such optimal experimental conditions may be hindered by, for example, the active regulation of pigment synthesis - processes that could be overcome by cellular engineering. Using our model of a cell without long-term adaption, we may identify optimal conditions to aim for in cell engineering and experimentation. By simulating a target compound consuming the amount of ATP, NADPH and reduced Ferredoxin (Fd) necessary to synthesize the target compound from carbon fixation, we tried to estimate the maximum production potential without limitation by the CBB. We found that the simulated production of all three compounds was highest under red light illumination (624 nm). Sucrose production was saturated at intermediate light and even showed slight inhibition under high light, while the simulated isoprene production, requiring reduced Fd and a lower amount of ATP, showed the highest requirement for light (no saturation at 1600 µmol(photons) m^*−*2^ s^*−*1^). Thus, the composition of energy equivalents seems to determine the optimal lighting conditions. NADPH production in particular seemed to follow a light saturation curve with maximum around 1600 µmol(photons) m^*−*2^ s^*−*1^ For the purpose of whole-cell biocatalysis, NADPH is often the only required cofactor for the reaction, while the generation of ATP and biomass are secondary. Studies have attempted to optimise NADPH regeneration through inhibition of the CBB, deletion of flavodiiron proteins, or introducing additional heterologous sinks for ATP, while at the same time trying to avoid oxidative stress [93, 94, 95]. Our simulations suggest that a switch in light color towards monochromatic red light may be a viable strategy to improve catalysis by matching the NADPH-focused demand of the sink reaction with an equally biased source reaction.

These results again support the need to test and optimize light conditions for each application on its own, as the stoichiometry of the desired process changes light requirements. Recently, two-phase processes have been used to increase titers in cyanobacterial biotechnology, arresting growth to direct all carbon towards a product [96]. Our model suggests that as a part of this process, changes in light color could be used to intentionally create imbalances in metabolism and direct flux to the desired product according to the energetic needs of the particular pathway.

To address the limitations of current model, it is imperative to critically evaluate its underlying assumptions and identify key areas for improvement. For instance, withe the current version of the model we cannot predict the long-term cellular adaption governed by many photoreceptors [97, 98]. For each simulation, we assume fixed pigment composition and light absorption capacity, thus, analysing a given cell state. Relevant cellular adaptions can, however, be used as new inputs according to experimental data. Also, rhodopsin photoreceptors can perform light-driven ion transport and, if found photosynthetically relevant, would be a useful addition to the model [99, 100]. Next, although our model considers the CBB as the main sink for energy equivalents, reactions downstream of the CBB, such as glycogen production [101]), could pose additional significant sinks depending on the cell’s metabolic state, necessitating further refinement of our model to accurately capture these dynamics. Additionally, further improvements of the currently significantly simplified CCM (Fig. S2) and photorespiratory salvage functions could be beneficial, also due to the engineering efforts in building pyrenoid-based CO_2_-concentrating mechanisms *in-planta* [102]. Photodamage may be a necessary addition to the model when considering high-light conditions, specifically PSII photoinhibition and the Mehler reaction [103] see i.e. [30]). Finally, our model follows the dynamic change in the lumenal and cytoplasmic pH but is lacking the full description of *pmf*. An envisaged step of further development will be the integration of the membrane potential ΔΨ into the model and simulation of ion movement, as presented in several mathematical models for plants [18, 104]. It would be moreover interesting to include the spatial component into the model, accounting for the dynamics of thylakoid membranes, as revealed by [105]. Thanks to our computational implementation of the model using the package `modelbase` [51], the model is highly modular, and the addition of new pathways or the integration of other published models (e.g. a recent CBB model [106]) should not constitute a technical challenge.

In conclusion, the development of our first-generation computational model for simulating photosynthetic dynamics represents a significant advancement in our comprehension of cyanobacteria-specific photosynthetic electron flow. While acknowledging its imperfections, our model has proven to be a versatile tool with a wide range of applications, spanning from fundamental research endeavors aimed at unravelling the complexities of photosynthesis to practical efforts focused on biotechnological optimization. Through a comprehensive presentation of our results, we have demonstrated the model’s capacity to elucidate core principles underlying photosynthetic processes, test existing hypotheses, and offer valuable insights on the photosynthetic control under various light spectra. With further development and integration of experimental data, we hope to provide a reference kinetic model of cyanobacteria photosynthesis.

## Acknowledgements

We would like to thank Ilka Axmann and Marion Eisenhut for the initial conversations on the physiology of cyanobacteria that motivated the construction of this detailed model, and David Fuente for the discussion on the cyanobacterial absorption and pigment compositions.

## Authors contribution

TP, EK developed the computational model; TP, EK and AN performed simulations and visualizations; TZ performed validation experiments and performed data curation; OE, AM, JC, GB acquired funding; TP, EK, AN, and AM wrote the original draft; TP and AM reviewed and edited the corrected manuscript; AM conceptualized, initiated and supervised the project; all authors were involved in discussions on methodology and investigation and made a substantial, direct, and intellectual contribution to the work and approved it for publication.

