## Supplemental Materials for "Shedding light on blue-green photosynthesis: A wavelength-dependent mathematical model of photosynthesis in *Synechocystis* sp. PCC 6803"

### Supplemental Materials: Shedding light on blue-green photosynthesis

### S1 Model structure

Our model consists of a system of 17 coupled Ordinary Differential Equations (ODEs), 27 reaction rates, and 95 parameters, including measured midpoint potentials, compound concentrations, absorption spectra, and physical constants. We have been using integration methods from the modelbase package [107] to solve the system with the initial values summarised in Table S1.

Table S1: **Initial conditions.** Initial values were set at the beginning of a simulation. For the calculations, see the GitLab file *calculate_parameters_restruct.py*.

| Parameter | Value | Description | Source value | Source |
|---|---|---|---|---|
| PSII | 1.232 [mmol mol(Chl)^-1] | initial concentration of unquenched PSII | estimated from cell being in state 2 in darkness | estimated |
| O2 | 55.402 [mmol mol(Chl)^-1] | concentration of oxygen in the cell | 230 [umol l^-1] concentration of oxygen in air saturated water: 230 uM | [108] |
| PC_ox | 0.157 [mmol mol(Chl)^-1] | initial concentration of oxidized plastocyanin (aerobic) | 0.1 [unitless] fraction of oxidized plastocyanin (aerobic) | [109] |
| Fd_ox | 3.324 [mmol mol(Chl)^-1] | initial concentration of oxidized ferredoxin (aerobic) | 0.9 [unitless] fraction of oxidized ferredoxin (aerobic) | [109] |
| NADPH | 20.104 [mmol mol(Chl)^-1] | initial concentration of NADPH | 0.75 [unitless] fraction of reduced NADPH | [53] |
| NADH | 3.574 [mmol mol(Chl)^-1] | initial concentration of NADH | 0.32 [unitless] approximate fraction of reduced NADH | [110] |
| ATP | 172.057 [mmol mol(Chl)^-1] | initial concentration of ATP | 400 [pmol (10^8 cells)^-1] concentration of ATP | [111] |
| PG | 0.894 [mmol mol(Chl)^-1] | initial concentration of (2-phospho) glycolate | concentration below 1e-6 [umol ug(Chl)^-1], the estimated detection limit of the method used by Huege (2011) | [112] |
| succinate | 2.000 [mmol mol(Chl)^-1] | initial concentration of succinate | estimated | estimated |
| fumarate | 2.000 [mmol mol(Chl)^-1] | initial concentration of fumarate | estimated | estimated |
| 3PGA | 2.000e+03 [mmol mol(Chl)^-1] | initial concentration of 3-phosphoglycerate (including all other sugars) | manually fitted to be non-limiting for RuBisCO oxygenation and respiration | manually fitted |
| CO2 | 3.103 [mmol mol(Chl)^-1] | concentration of $CO_2$ in the cell without activity of the CCM | 322e-4 [mol l^-1 atm^-1] solubility of CO2 in 25 °C water with ~10 ‰ Cl^- ions | [113] |
| CBBa | 0.000e+00 [unitless] | initial redox-regulated activity of the CBB | CBB should be inactive in the dark to avoid futile cycles | estimated |
| Hi | 0.217 [mmol mol(Chl)^-1] | initial concentration of lumenal protons | pH 5 in 10^4 uE cm^-1 s^-1 light | [52] |
| Ho | 6.932e-03 [mmol mol(Chl)^-1] | initial concentration of cytoplasmic protons | pH 7.5 in 10^4 uE cm^-1 s^-1 light | [52] |
| Q_ox | 7.202 [mmol mol(Chl)^-1] | concentration of oxidized plastoquinone | 0.446 [unitless] fraction of PHOTOACTIVE plastoquinone reduced in 40 umol m^-2 s^-1 light | [54] |
| OCP | 0.000e+00 [unitless] | initial activity of OCP | OCP quenching is relaxed in the dark | estimated |

#### S1.1 A complete summary of the modelled reactions

The model's main body consists of these four electron transport pathways:

- LET: Electrons originating from PSII and follow the redox potential from PQ via $Cb_6f$ to PC to PSI. In a second light reaction, electrons are excited again, transferred to Fd, and afterward to $NADP^+$ by the FNR.
- RET: Alternatively, electrons could originate from carbon compound respiration. These electrons can be transferred **a)** from the TCA intermediate fumarate to PQ via succinate dehydrogenase; **b)** from NADH to PQ via NADH dehydrogenase-like complex type-2; or **c)** backwards from NADPH to Fd via FNR. These electrons typically leave the ETC through a terminal oxidase.
- CET: From Fd, the electrons can also travel back to PQ via the NAD(P)H Dehydrogenase-like complex 1. The electrons, therefore, cycle around PSI.
- AEF: Lastly, electrons originating from PSII can also be transferred to $O_2$ by a terminal oxidase (bb-type, $aa_3$-type, or Flv) without affecting the NADPH redox state.

Protons pumped during electron transport drive the ATP generation through ATP synthase or leak across the membrane. The CBB's carbon fixation is the leading consumer of ATP and NADPH. Another sink is the consumption of RuBisCO oxygenation side products through PR salvage. Additionally, consuming reactions for ATP and NADH simulate the remaining metabolic demand.

Both FNR and CBB are activated by reduced Fd while PQ induces PSII quenching. Lastly, $O_2$ diffuses across the cell membrane while the CCM actively imports $CO_2$. For a complete list of reactions, see Table S2.

Table S2: **The model reactions with stoichiometry.** The reactions are grouped according to electron transport pathways or similar functions. Parentheses group logically connected compounds, while square brackets mark compounds that are necessary for mass balance but are not part of the model.

| Description | Reaction | Name in code |
|---|---|---|
| | Linear Electron Transport | |
| Light reaction of PSII | $(PQ + 2\,H_o^+)\,[+H_2O] \longleftrightarrow PQH_2 + (2\,H_i^+ + 0.5\,O_2)$ | PS2 |
| Cytochrome $b_6f$ complex | $PQH_2 + 2\,PC_{ox} + 2\,H_o^+ \longleftrightarrow (PQ + 2\,H_i^+) + 2\,PC_{red} + 2\,H_i^+$ | b6f |
| Light reaction of PSI | $PC_{red} + Fd_{ox} \longleftrightarrow PC_{ox} + Fd_{red}$ | PS1 |
| Ferredoxin-$NADP^+$ Reductase | $(NADP^+ + H_o^+) + 2\,Fd_{red} \longleftrightarrow NADPH + 2\,Fd_{ox}$ | FNR |
| | Respiratory Electron Transport | |
| Respiration of sugar compounds with multiple pathways | $3PGA + 7.402\,e\text{-}02\,fumarate + (0.567\,ADP + [0.567\,P_i]) + 2.237\,NADP^+ + 2.689\,NAD^+ \longleftrightarrow$ <br> $3\,CO_2 + 7.402\,e\text{-}02\,succinate + 0.567\,ATP + (2.237\,NADPH + 2.237\,H_o^+) + (2.689\,NADH + 2.689\,H_o^+)$ | Respiration |
| Succinate dehydrogenase | $(PQ + 2\,H_o^+) + succinate \longleftrightarrow PQH_2 + (fumarate + 2\,H_o^+)$ | SDH |
| NADH dehydrogenase-like complex type-2 | $NADH + (PQ + 2\,H_o^+) \longleftrightarrow (NAD^+ + H_o^+) + PQH_2$ | NDH |
| | Cyclic Electron Transport | |
| NADH dehydrogenase-like complex type-1 | $2\,Fd_{red} + (PQ + 2\,H_o^+) + H_o^+ \longleftrightarrow 2\,Fd_{ox} + PQH_2 + H_i^+$ | NQ |
| | Alternate Electron Transport | |
| bd-type terminal oxidase | $2\,PQH_2 + (O_2 + 4\,H_o^+) \longrightarrow (2\,PQ + 4\,H_i^+)\,[+2\,H_2O]$ | bd |
| aa3-type terminal oxidase (active proton pump) | $4\,PC_{red} + (O_2 + 5\,H_o^+) \longrightarrow 4\,PC_{ox}\,[+2\,H_2O] + H_i^+$ | aa |
| Flavodiiron protein dimer 1/3 | $4\,Fd_{red} + (O_2 + 4\,H_o^+) \longrightarrow 4\,Fd_{ox}\,[+2\,H_2O]$ | Flv |
| | ATP synthase and proton leak | |
| F0F1 ATPase | $(ADP + [P_i]) + HPR \cdot H_i^+ \longleftrightarrow ATP + HPR \cdot H_o^+$ | ATPsynthase |
| Proton leakage across the thylakoid membrane | $H_i^+ \longrightarrow H_o^+$ | Pass |
| | Calvin-Benson-Bassham cycle and Photorespiration | |
| RuBisCO carboxylation and Calvin-Benson-Bassham cycle | $(3\,CO_2 + 10\,H_o^+) + 8\,ATP + 5\,NADPH \longrightarrow 3PGA + 8\,ADP + (5\,NADP^+ + 5\,H_o^+)$ | CBB |
| RuBisCO oxygenation (includes steps of the CBB) | $(2 \cdot 3PGA + 3\,O_2 + 10\,H_o^+) + 8\,ATP + 5\,NADPH \longrightarrow 3 \cdot PG + 8\,ADP + (5\,NADP^+ + 5\,H_o^+)$ | Oxy |
| Photorespiratory salvage pathway (multiple mechanisms) | $2 \cdot PG + ATP + NADPH + 2\,NAD^+ \longleftrightarrow (CO_2 + 3PGA) + (ADP\,[+P_i]) + (NADP^+ + H_o^+) + 2\,NADH$ | PRsalv |
| | Consuming reactions | |
| Cellular metabolic consumption of ATP | $ATP \longrightarrow ADP\,[+P_i]$ | ATPconsumption |
| Cellular metabolic consumption of NADH | $NADH + (+2\,H_o^+\,[+C]) \longrightarrow (NAD^+ + H_o^+)\,[+CH_2]$ | NADHconsumption |
| | Regulatory reactions | |
| Fd-dependent activation of the CBB | $CBB_{inactive} \longleftrightarrow CBB_{active}$ | CBBactivation |
| PSII-internal quenching (mechanism BASED ON [67]) | $PSII_{unquenched} \longleftrightarrow PSII_{quenched}$ | PSIIquench |
| | Gas exchange | |
| $O_2$ diffusion out of the cell | $O_2 \longleftrightarrow$ | O2out |
| CCM driven $CO_2$ transport into the cell | $\longleftrightarrow CO_2$ | CCM |

#### S1.2 Simplified irreversible mass action kinetics

Kinetic descriptions that don't follow MA or Equation 1 are given below. For simplifying the reversible MA law [57] we followed the approach of Noor *et al.* [58] by separating the kinetic and thermodynamic terms within the rate law. Exemplary for the forward direction ($\Delta_{\mathrm{r}}\mathrm{G}^{'} < 0$):

$$\mathrm{v} = \mathrm{k} \cdot \left( \prod c_{S_i}^{n_i} - \frac{\prod c_{P_j}^{m_j}}{K_{eq}} \right) = \overbrace{\mathrm{k} \cdot \prod c_{S_i}^{n_i}}^{\text{kinetic}} \cdot \overbrace{\left( 1 - \frac{\prod c_{P_j}^{m_j} / \prod c_{S_i}^{n_i}}{K_{eq}} \right)}^{\text{thermodynamic ("}\gamma\text{")}} \tag{S1}$$

with $K_{eq} = \exp(-\Delta_r G^{'0}/\mathrm{RT})$ and $\Delta_r G^{'} = \Delta_r G^{'0} + RT \cdot \ln(\prod c_{P_j}^{m_j} / \prod c_{S_i}^{n_i})$ this can be written as

$$= \mathrm{k} \cdot \prod c_{S_i}^{n_i} \cdot \Big( 1 - \exp(\Delta_r G^{'}/\mathrm{RT}) \Big) \tag{S2}$$

and simplifying only the kinetic term

$$v \approx \mathrm{k}^{+} \cdot \prod c_{S_i} \cdot \Big( 1 - \exp(\Delta_r G^{'}/\mathrm{RT}) \Big) \tag{S3}$$

We did not alter the thermodynamic term to maintain $\mathrm{v} = 0$ at the MA-derived equilibrium concentrations. We proceeded similarly for the reverse reaction ($\Delta_{\mathrm{r}}\mathrm{G}^{'} > 0$), factoring out $\prod c_{P_j}^{m_j}/\mathrm{K_{eq}}$:

$$\mathrm{v} = \mathrm{k} \cdot \frac{\prod c_{P_j}^{m_j}}{K_{eq}} \cdot \left( \frac{K_{eq}}{\prod c_{P_j}^{m_j} / \prod c_{S_i}^{n_i}} - 1 \right) \tag{S4}$$

$$\approx \mathrm{k}^{-} \cdot \frac{\prod c_{P_j}}{K_{eq}} \cdot \Big( \exp(-\Delta_r G^{'}/\mathrm{RT}) - 1 \Big) \tag{S5}$$

By simplifying in S3 and S5 we approximate

$$k \cdot \prod c_{S_i}^{n_i} \approx k^{+} \cdot \prod c_{S_i}^{n_i} / \prod c_{S_i}^{n_i - 1} \quad \text{and} \quad k \cdot \prod c_{P_j}^{m_j} \approx k^{-} \cdot \prod c_{P_j}^{m_j} / \prod c_{P_j}^{m_j - 1} \tag{S6}$$

which means for the relevant case of non-zero rates ($\prod c_{S_i}^{n_i} > 0$ and $\prod c_{P_j}^{m_j} > 0$)

$$k \approx \frac{\mathrm{k}^{+}}{\prod c_{S_i}^{n_i - 1}} \approx \frac{\mathrm{k}^{-}}{\prod c_{P_j}^{m_j - 1}} \tag{S7}$$

For any $n_i > 1$ or $m_j > 1$, the two denominators may not be equal and thus

$$\mathrm{k}^{+} \neq \mathrm{k}^{-} \tag{S8}$$

#### S1.3 Description of photosystems

We model the photosystems using QSS equation systems similar to our previous models [47, 48] representing PSII as a four-state model and PSI as three-state model.

The four-state model of PSII consists of the open and closed reaction center states ($B_0$, $B_2$) as well as their respective excited states ($B_1$, $B_3$) (Fig. S1). The PSII excitation rate constant $k_{LII}$ is calculated from $E_{PSI}$ in Equation 2 (in µmol(photons) mg(chl)$^{-1}$) by multiplication with the molar mass of chlorophyll $M_{Chl}$ and dividing by the PSII concentration:

$$k_{LII} = E_{PSI} \cdot M_{Chl} \cdot \frac{1}{c_{II}} \tag{S9}$$

The resulting excited states can be quenched by photochemistry (rate constant ($k$): $k_2$, only $B_1$), as heat ($k$: $k_H$) or fluorescence ($k$: $k_F$). PSII with closed reaction centers can also reversibly reduce PQ ($k$: $k_{PQred}$). The QSS assumption together with the conservation of PSII complexes results in the equation system:

$$\frac{dB_0}{dt} = -(k_{LII} + \frac{k_{PQred} PQ_{red}}{K_{eq,PQred}}) \cdot B_0 + (k_H + k_F) \cdot B_1 + k_{PQred} PQ_{ox} \cdot B_2 = 0 \tag{S10}$$

$$\frac{dB_1}{dt} = k_{LII} \cdot B_0 - (k_H + k_F + k_2) \cdot B_1 = 0 \tag{S11}$$

$$\frac{dB_3}{dt} = k_{LII} \cdot B_2 - (k_H + k_F) \cdot B_3 = 0 \tag{S12}$$

$$B_0 + B_1 + B_2 + B_3 = PSII_{tot} \tag{S13}$$

Note that state transitions affect $k_H$ in the PSII quenching model (see Equation S35). We defined the PSII rate $v_{PSII}$ as

$$v_{PSII} = 0.5 \cdot k_2 \cdot B_1 \tag{S14}$$

since two $B_1 \rightarrow B_2$ reactions have to occur for a full PQ reduction.

PSI was described similarly using a three-state QSS system. The open state $Y_0$ is excited to $Y_1$ ($k$: $k_{L1}$), which can donate an electron to Fd ($k$: $k_{Fdred}$). The resulting oxidized PSI ($Y_2$) is reduced by PC ($k$: $k_{PCox}$) to return to the open state. To represent PSI fluorescence, we have added a slow relaxation of $Y_1$ by fluorescence ($k$: $k_{F1}$). The resulting equation system

$$\frac{dY_0}{dt} = -(\frac{k_{PCox}PC_{ox}}{K_{eq,PCP700}} + k_{LI}) \cdot Y_0 + k_{F1} \cdot Y_1 + k_{PCox}PC_{red} \cdot Y_2 = 0 \tag{S15}$$

$$\frac{dY_1}{dt} = k_{LI} \cdot Y_0 - (k_{F1} + k_{Fdred}Fd_{ox}) \cdot Y_1 + \frac{k_{Fdred}Fd_{red}}{K_{eq,FAFd}} \cdot Y_2 = 0 \tag{S16}$$

$$Y_0 + Y_1 + Y_2 = PSI_{tot} \tag{S17}$$

is also symbolically solved at every integration step and used to determine the PSI reaction rate $v_{PSI}$ as the net rate of excitations

$$v_{PSI} = k_{L1} \cdot Y_0 - k_{F1} \cdot Y_1 \tag{S18}$$

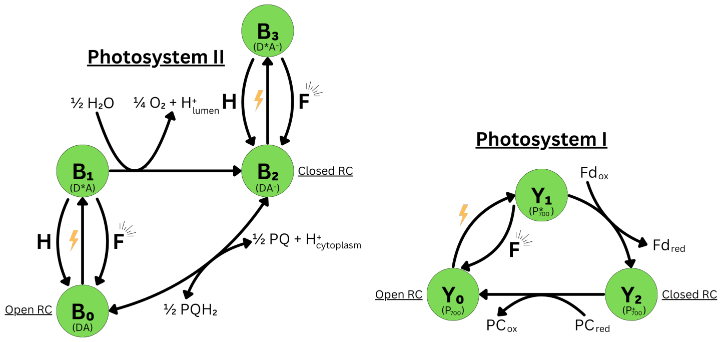

Fig. S1: **Schematic of the photosystem's modelled internal processes.** a) Photosystem II. The open reaction centers (RC) $B_0$ are excited by light (yellow bolt). The excited state $B_1$ can relax to $B_0$ by heat ($H$) and fluorescence ($F$) emission or perform photochemistry. The latter promotes the RC to the closed state $B_2$ and extracts one electron from water. Excitation of $B_2$ can only be quenched as $H$ or $F$. Lastly, $B_2$ can reduce Plastoquinone (PQ) and enter the open state $B_0$ again. Parentheses show the assumed state of the special pair chlorophyll $P_{680}$ ($D$) and electron acceptor plastoquinone A ($A$): excited (*) and reduced($^-$). b) Photosystem I. Light excites the open reaction centers $Y_0$. The excited $Y_1$ state can perform photochemistry by reducing Ferredoxin (Fd) and becoming oxidized to $Y_2$. We also consider a minor relaxation of $Y_1$ to $Y_0$ through $F$. The oxidized $Y_2$ is reduced by Plastocyanine (PC). Parentheses show the assumed state of the reaction center $P_{700}$.

#### S1.4 Exemplary Gibbs free energy calculation

Here, we explain the calculation of Gibbs free energies for reaction kinetics on the example of FNR. First, we calculate the standard Gibbs free energy ($\Delta_r G^{'0}$) following Ebenhöh *et al.* [47]. We split the model reaction into redox pair reactions with electrons $e^-$ and protons $H^+$:

$$\mathrm{Ox} + \mathrm{a\,e^-} + \mathrm{b\,H^+} \longleftrightarrow \mathrm{Red} \quad \text{with } \mathrm{a, b} \in \mathbb{N} \tag{S19}$$

If the reaction doesn't involve free elections, the sub reaction's $\Delta_r G^{'0}$ is calculated as [47]:

$$\Delta_r G^{'0} = -\mathrm{a} \cdot \mathrm{F} \cdot \mathrm{E_0} + \mathrm{b} \cdot ln(10)RT \cdot \mathrm{pH} \tag{S20}$$

With the Faraday constant $F$, the sub reaction's standard electrode potential $E_0$, the ideal gas constant $R$, and temperature $T$. The total $\Delta_r G^{'0}$ is then the stoichiometric sum of the sub reaction $\Delta_r G^{'0}$. We can decompose the FNR reaction (excluding the NADPH-associated proton)

$$2\,\mathrm{Fd_{red}} + \mathrm{NADP^+} + \mathrm{H^+} \longleftrightarrow 2\,\mathrm{Fd_{ox}} + \mathrm{NADPH} \tag{S21}$$

into the sub reactions

$$\mathrm{Fd_{ox}} + \mathrm{e^-} \longleftrightarrow \mathrm{Fd_{red}} \tag{S22}$$

$$\mathrm{NADP^+} + 2\,\mathrm{e^-} + \mathrm{H^+} \longleftrightarrow \mathrm{NADPH} \tag{S23}$$

with stoichiometry $(S21) = -2\cdot(S22)+1\cdot(S23)$. We then calculate the $\Delta_r G^{'0}$ according to Equation S20:

$$\Delta_r G^{'0}_{Fd} = -1 \cdot \mathrm{F} \cdot \mathrm{E}_0 \tag{S24}$$

$$\Delta_r G^{'0}_{NADP} = -2 \cdot \mathrm{F} \cdot \mathrm{E}_0 + 1 \cdot ln(10)RT \cdot \mathrm{pH} \tag{S25}$$

$$\Delta_r G^{'0}_{FNR} = -2 \cdot \Delta_r G^{'0}_{Fd} + 1 \cdot \Delta_r G^{'0}_{NADP} \tag{S26}$$

At each time step we now calculate Gibbs free energy $\Delta_r G^{'}$ using the reactant concentrations:

$$\Delta_r G^{'} = \Delta_r G^{'0} + RT \cdot \ln(\textstyle\prod c_{P_j}^{m_j} / \prod c_{S_i}^{n_i}) \tag{S27}$$

$$\Delta_r G^{'}_{FNR} = \Delta_r G^{'0}_{FNR} \cdot \frac{\mathrm{Fd_{red}^2} \cdot \mathrm{NADP^+}}{\mathrm{Fd_{ox}^2} \cdot \mathrm{NADPH}} \tag{S28}$$

With $\Delta_r G^{'}$ and $\Delta_r G^{'0}$ we can then calculate the reaction rate in Equation 1.

#### S1.5 Calculating pigment association

In addition to PBS, we calculate the fraction of light absorbed by chlorophyll and beta-carotene passed to the photosystems. For chlorophyll, the relationship is given as the ratio of their bound chlorophyll:

$$p_{Chl,I} = 1 - p_{Chl,II} = \frac{3c_I \cdot n_I^{Chl}}{3c_I \cdot n_I^{Chl} + 2c_{II} \cdot n_{II}^{Chl}} \tag{S29}$$

Similarly, we calculate the relative amounts of beta-carotene bound to either photosystem. However, as described by Fuente *et al.* [59], beta-carotene is also present outside of the photosystems. The photosystem stoichiometry requires a certain ratio of beta-carotene to chlorophyll.

$$f_{Chl:Car,stoich} = \frac{3c_I \cdot \frac{n_I^{Chl}}{n_I^{Car}} + 2c_{II} \cdot \frac{n_{II}^{Chl}}{n_{II}^{Car}}}{3c_I + 2c_{II}} \tag{S30}$$

We assume that any beta-carotene present in the cell above that ratio is not associated with the photosystems. Therefore we calculate the fraction beta-carotene absorption passed to photosystems $f_{Car}$ as:

$$f_{Car} = \frac{f_{Chl:Car,stoich}}{f_{Chl:Car,meas}} \tag{S31}$$

$$p_{Chl,I} = \frac{3c_I \cdot n_I^{Car}}{3c_I \cdot n_I^{Car} + 2c_{II} \cdot n_{II}^{Car}} \cdot f_{Car} \tag{S32}$$

$$p_{Chl,II} = \frac{2c_{II} \cdot n_{II}^{Car}}{3c_I \cdot n_I^{Car} + 2c_{II} \cdot n_{II}^{Car}} \cdot f_{Car} \tag{S33}$$

where $f_{Chl:Car,meas}$ is the measured molar ratio of beta carotene to chlorophyll.

#### S1.6 Possible mechanisms of state transitions

We have implemented four possible mechanisms of state transitions.

---

**PSII-quenching Model** (default): In the presence of $PQ_{red}$, PSII reversibly enters a quenched state $PSII_q$. This increases the rate of overall excitation quenching as heat ($k_H$).

$$\frac{dPSII_q}{dt} = k_{Quench} \cdot (1 - PSII_q) \cdot PQ_{red} - \frac{k_{Unquench} \cdot PSII_q \cdot PQ_{red}^{\mathrm{n_{Unquench}}}}{\mathrm{K_{M,Unquench}^{n_{Unquench}}} + PQ_{red}^{\mathrm{n_{Unquench}}}} \tag{S34}$$

$$k_H = k_{H0} + k_{Hst} \cdot (PSII_q/\mathrm{PSII_{tot}}) \tag{S35}$$

---

**Spillover**: State 2 transition increases the fraction $spill(\leq spill_{max})$ of PSII excitations ($E_{PSII}$) passed to PSI.

$$\frac{dspill}{dt} = k_{spill} \cdot (spill_{max} - spill) \cdot PQ_{red} - k_{unspill} \cdot spill \cdot PQ_{ox} \tag{S36}$$
$$E_{PSII,spill} = E_{PSII} \cdot (1 - spill) \tag{S37}$$
$$E_{PSI,spill} = E_{PSI} + E_{PSII} \cdot spill \tag{S38}$$

---

**PBS-Mobile**: In state 2 transition, PBS move with rate $v_{toPSI}$ from PSII ($PBS_{II} \geq PBS_{II,min}$) to PSI ($PBS_{I} \geq PBS_{I,min}$).

$$v_{toPSI} = k_{toPSI} \cdot (PBS_{II} - PBS_{II,min}) \cdot PQ_{red} - k_{toPSII} \cdot (PBS_{I} - PBS_{I,min}) \cdot PQ_{ox} \tag{S39}$$
$$\frac{dPBS_{I}}{dt} = -\frac{dPBS_{II}}{dt} = v_{toPSI} \tag{S40}$$

---

**PBS-Detachment**: In state 1 transition PBS detach equally from both photosystems (to $PBS_{free} \leq PBS_{free,max}$) with rate $v_{detach}$.

$$v_{detach} = k_{detach} \cdot (PBS_{free,max} - PBS_{free}) \cdot PQ_{ox} - k_{attach} \cdot PBS_{free} \cdot PQ_{red} \tag{S41}$$
$$\frac{dPBS_{free}}{dt} = -0.5\frac{dPBS_{II}}{dt} = -0.5\frac{dPBS_{I}}{dt} \tag{S42}$$

---

#### S1.7 Estimating pathway fluxes

We estimate the flux through the four main electron pathways from reactions that are preferably unique to the pathway. Because electrons from respiration also enter the PETC, we scale the fluxes of LET and AEF, pathways that are defined beginning with PSII, with the fraction of PSII-derived electron flux

$$f_{PSII}^{influx} = \frac{2 \cdot vPSII}{2 \cdot vPSII + 2 \cdot vSDH + 2 \cdot vNDH2} \tag{S43}$$

All reaction fluxes are scaled by the number of involved electrons

$$vLET = 2 \cdot vFNR \cdot f_{PSII}^{influx} \tag{S44}$$
$$vCET = 2 \cdot vNDH1 \tag{S45}$$
$$vRET = 2 \cdot vSDH + 2 \cdot vNDH2 \tag{S46}$$
$$vAEF = (4 \cdot vFlv + 4 \cdot vCyd + 4 \cdot vCOX) \cdot f_{PSII}^{influx} \tag{S47}$$

We do not calculate these electron fluxes under very low light intensities ($< 10\,\mu mol(photons)\,m^{-2}\,s^{-1}$) where FNR flux can be reversed due to high NADPH concentrations stemming from respiration.

#### S1.8 Estimating fluorescence parameters and heat quenching

For the evaluation of Fig. S7 we calculated the fluorescence parameters Non-Photochemical Quenching (NPQ) and the quantum yield of PSII (Y(II)). Both parameters are estimated during a PAM-SP experiment from the dark-adapted maximal fluorescence ($F_m$), the maximal fluorescence in the light ($F'_m$), and the steady-state fluorescence (F) as [22]

$$NPQ = \frac{F_m - F'_m}{F'_m} \tag{S48}$$
$$Y(II) = \frac{F'_m - F}{F'_m} \tag{S49}$$

Since these quantities are defined on the basis of PSII chlorophyll fluorescence, we only consider the PSII fluorescence component $F_{PSII}$ in these calculations. We simulate a saturation pulse (600 ms, 633 nm light, 15 000 $\mu mol(photons)\,m^{-2}\,s^{-1}$) after oxidation of the PQ pool by 300 s of 700 nm monochromatic far-red light ($F_m$) or following the steady-state simulation at a particular light intensity ($F'_m$).

#### S1.9 RuBisCO reactions

The cyanobacterial CBB is known to be redox-regulated, with thioredoxin being a possible regulator [114, 115]. As thioredoxin is reduced from the Fd pool, we defined a Fd-dependent CBB activity regulator $CBB_a$ using Hill kinetics

$$\frac{dCBB_a}{dt} = \mathrm{k_{CBBa}} \cdot Fd_{red}^{n_{CBB}} / (Fd_{red}^{n_{CBB}} + K_{Hill,CBB}^{n_{CBB}}) \tag{S50}$$

Additionally, we modelled the CBB's dependency on ATP and NADPH with MM kinetics. As $CO_2$ and $O_2$ compete for binding to RuBisCO, we modelled their influence as competitive inhibition.

$$v_{CBB} = v_{CBB,max} \cdot CBB_a \cdot \frac{ATP}{ATP + K_{M,ATP}} \cdot \frac{NADPH}{NADPH + K_{M,NADPH}} \cdot \frac{CO_2}{K_{M,CO_2}(1 + {}^{O_2}/_{K_{I,O_2}})} \tag{S51}$$

We model the $O_2$-dependent RuBisCO oxygenation reaction similarly. Since the oxygenation consumes ribulose-1,5-bisphosphate, the lumped stoichiometry requires CBB regeneration reactions. Therefore, we also impose the redox, ATP, and NADPH constraints. Additionally, we limit oxygenation at low concentration of organic carbon, i.e. 3PGA.

$$v_{Oxy} = v_{Oxy,max} \cdot CBB_a \cdot \frac{ATP}{ATP + K_{M,ATP}} \cdot \frac{NADPH}{NADPH + K_{M,NADPH}} \cdot \frac{O_2}{K_{M,O_2}(1 + {}^{CO_2}/_{K_{I,CO_2}})} \cdot \frac{3PGA}{3PGA + K_{M,3PGA}} \tag{S52}$$

#### S1.10 Flavodiiron proteins

Measurements of $O_2$ reduction have shown that Flv only reduces oxygen under high light intensities (Fig. S4). We replicate this behavior using Hill dependency on the likely limiting substrate, Fd:

$$v_{Flv} = \mathrm{k_{Flv}} \cdot O_2 \cdot H_O^+ \cdot Fd_{red}^{n_{Flv}} / (Fd_{red}^{n_{Flv}} + K_{Hill,Flv}^{n_{Flv}}) \tag{S53}$$

With $H_O^+$ being cytoplasmic protons used for $O_2$ reduction to water.

#### S1.11 The orange carotenoid protein

The Orange Carotenoid Protein (OCP) is known to be activated by its carotenoid cofactor absorbing light, which we calculate using the OCP absorption spectrum vector $a_{OCP}$ [116]. We represent the inactivation of OCP by the fluorescence recovery protein through a MA kinetic. A maximum value $OCP_{max}$ limits the amount of PBS excitations quenched.

$$\frac{dOCP}{dt} = (OCP_{max} - OCP) \cdot k_{OCPactivation} \cdot \mathrm{simpson}(\mathrm{diag}(I) \cdot a_{OCP}) \cdot lcf - OCP \cdot k_{OCPdeactivation} \tag{S54}$$

#### S1.12 Cellular import of $CO_2$

Cyanobacteria increase the carbon concentration around RuBisCO actively by the factor 100 - 1000 through the CCM and the construction of carboxysomes [117]. We model this by increasing the intracellular $CO_2$ partial pressure $p_{CO_2,in}$ by the factor $f_{CO_2,in}$:

$$p_{CO_2,in} = p_{CO_2,in} \cdot f_{CO_2,in} \tag{S55}$$

The concentration of $CO_2$ that can dissolve ($CO_{2,sol}$) in water is given by Henry's law [118]:

$$CO_{2,sol} = {}^{p_{CO_2,in}}/_{K_{CO_2}} \tag{S56}$$

$$\ln(K_{CO_2}) = -58.0931 + 90.5069\frac{100}{T} + 22.2940 \cdot \ln\left(\frac{T}{100}\right) + S\left(0.027766 - 0.025888 * \frac{T}{100} + 0.0050578(\frac{T}{100})^2\right) \tag{S57}$$

where the Henry constant $K_{CO_2}$ depends on the temperature $T$ and (intracellular) salinity $S$. Furthermore, $CO_2$ dissociates into ${HCO_3}^-$ and ${CO_3}^{2-}$ which are not substrates of RuBisCO. Therefore, we calculate the maximal usable $CO_2$ concentration $CO_{2,max}$ using the first dissociation constant $K_1$ [119]

$$CO_{2,max} = CO_{2,sol} \cdot \frac{H_O^+}{H_O^+ + K_1} \tag{S58}$$

$$pK_1 = -43.6977 - 0.0129037S + 1.364e - 4S^2 + \frac{2885.378}{T} + 7.045159 \cdot \ln(T) \tag{S59}$$

$$K_1 = 10^{-pK_1} \cdot \frac{1}{c_{Chl}} \tag{S60}$$

where $K_1$ is converted into $\mathrm{mmol\,mol(chl)}^{-1}$. We describe the dynamic intracellular $CO_2$ concentration in fast equilibrium with $CO_{2,max}$:

$$\frac{dCO_2}{dt} = k_{CCM}(CO_{2,max} - CO_2) \tag{S61}$$

### S2 Model parametrization

#### S2.1 Model parameters

A complete list of all parameters can be found in Table S3, Table S4 (strain-specific parameters) and Table S5 (for state transitions model).

The model depends on 95 parameters of which 37 were taken directly from literature (including 5 pigment absorption spectra), 9 rate parameters were estimated from direct or related literature rate measurements by determining an approximate rate through the pathway and dividing this rate by the assumed intracellular substrate concentrations, 6 parameters were physical quantities, 6 parameters were set according to the experimental irradiance, measuring light, O2 & CO2 condition, the concentration of cells, and temperature, 8 parameters were estimated from experimental data (PBS_PS2, PBS_PS1, PBS_free, PSItot, PSIItot, and pigment concentrations), 2 inhibition constants were estimated assuming they are equal to the Michaelis constants, the intracellular salinity S was assumed to be equal to seawater, intracellular buffering capacities of lumen and cytoplasm (bHo and bHi) were assumed constant [47], and 25 were fitted to different datasets or literature during model refinement:

- kUnquench, kQuench, KMUnquench, kOCPactivation, kOCPdeactivation, OCPmax, and PBS in fluo_influence were fitted to PAM-SP data [56] (Fig. 2a),
- parameter vNQ_max was fit to 30% CET measured in cyanobacteria [46] (Fig 1b),
- parameter lcf was fit to 15 electrons PSI-1 s-1 of LET under 300 µmol m-2 s-1 light [46] (Fig 1b)
- parameter kCBBactivation was fit to activate the CBB within ca. one minute [55],
- parameter KMFdred was fit to have a low CBB activity in the dark but high activity at irradiances where respiration is inhibited (at irradiance above ca. 10 µmols m-2 s-1) [25],
- parameters KHillFdred and nHillFdred were fit to induce fast Flv activation at ca. 50 µmols m-2 s-1 white irradiance [25],
- parameter fCin was fit to CO2 assimilation data [120],
- parameters k_respiration and k_aa were fit to ca. 20% PQ reduction in dark [54],
- parameter k_Q was fit to a level allowing for PQ oxidation under blue irradiance [41],
- parameter KMPGA was fit to prohibit Photorespiration under very low concentration of CBB intermediates (represented by 3PGA),
- parameters KMNQ_Qox and KMNQ_Fdred were fit to produce continuous CET flow under high PQ reduction near light saturation [46],
- parameter kATPsynth was fit to produce sufficient ATP for supplying the CBB,
- parameter k_F1 was fit to provide a low fluorescence yield to PSI,

- parameter kNADHconsumption was fit to provide reasonable NADH/NAD+ ratios in darkness and light, and
- kATPconsumption and k_pass were fit to provide low capacity flows dissipating ATP and proton gradients.

The final model has been validated against the newly obtained data (added to Fig. 2c, p. 9).

Table S3: **The model parameters with descriptions and sources.** Parameters with source "derived" are calculated from other parameters. See Table S1 for concentration parameters. For the parameter calculations, see the GitLab file *calculate_parameters_restruct.py*.

| Parameter | Value | Description | Source value | Source |
|---|---|---|---|---|
| | | Cell specific | | |
| V_cell | 5.600e-15 [l cell^-1] | synechocystis cell volume | 5.6e-15 [l cell^-1] synechocystis cell volume | [121] |
| n_chl | 1.400e+07 [cell^-1] | total chlorophyll content | 1.4e7 [cell^-1] total chlorophyll content | [122] |
| bHi | 100.000 [unitless] | buffering constant of the thylakoid lumen | estimated, like Ebenhöh (2014) | estimated |
| bHo | 1.111e+03 [unitless] | buffering constant of the cytoplasm, assumed to be 1/f_V_lumen times larger | bHi, f_V_lumen | estimated |
| f_V_lumen | 9.000e-02 [unitless] | thylakoid lumen fraction of the total cell volume | 0.09 [unitless] thylakoid lumen fraction of the total cell volume | [123] |
| V_cyt | 5.096e-15 [l cell^-1] | volume of the cytoplasm | f_V_lumen, V_cell | derived |
| V_lumen | 5.040e-16 [l cell^-1] | volume of the thylakoid lumen | f_V_lumen, V_cell | derived |
| cChl | 4.151e-03 [mol l^-1] | total molar concentration of chlorophyll | n_chl, V_cell, NA | derived |
| chl_cyt | 4.562e-03 [mol l^-1] | molar concentration of chlorophyll relative to the cytoplasmic volume | n_chl, V_cyt, NA | derived |
| chl_lumen | 4.613e-02 [mol l^-1] | molar concentration of chlorophyll relative to the lumen volume | n_chl, V_lumen, NA | derived |
| cf_lumen | 4.613e-05 [mol(Chl) ml^-1] | conversion factor for [mmol mol(Chl)^-1] -> [mol l^-1] for the thylakoid lumen | chl_lumen | derived |
| cf_cytoplasm | 4.562e-06 [mol(Chl) ml^-1] | conversion factor for [mmol mol(Chl)^-1] -> [mol l^-1] for the cytoplasm | chl_cyt | derived |
| fCin | 100.000 [unitless] | ratio of intracellular to external CO2 partial pressure with activity of the CCM | the CCM-increased intracellular bicarbonate concentration can exceed the external by 100 - 1000 times, manually fitted to Fig. S2 | [117], manually fitted |
| HPR | 4.667 [unitless] | number of protons (14) passing through the ATP synthase per ATP (3) synthesized | 14.0 protons per full rotation of ATP synthase | [124] |
| pigment_content | {chla: 1.000, beta_carotene: 0.176, allophycocyanin: 1.118 , phycocyanin: 6.765} [mg(Pigment) mg(Chla)^-1] | relative pigment concentrations in a synechocystis cell | | [56] |
| PBS_free | 9.000e-02 [unitless] | fraction of unbound PBS | measured PBS fluorescence at 77K | [56] |
| PBS_PS1 | 0.390 [unitless] | fraction of PBS bound to PSI | measured PBS fluorescence at 77K | [56] |
| PBS_PS2 | 0.510 [unitless] | fraction of PBS bound to PSII | measured PBS fluorescence at 77K | [56] |
| fluo_influence | {PS2: 1.000, PS1: 1.000, PBS: 1.250} [unitless] | factors multiplied to the calculated fluorescence (no effect at 1) | manually fitted to Fig. 2a | manually fitted |
| lcf | 0.500 [excitations photons^-1] | light conversion factor used for photosystem excitations, OCP activation, and PBS fluorescence | manually fitted to reproduce 15 electrons PSI^-1 s^-1 in Fig. 1b | manually fitted |
| | | Physical constants | | |
| F | 96.485 [C mmol^-1] | Faraday's constant | 9.6485e4 [C mol^-1] Faraday's constant | [125] |
| R | 8.300e-03 [J K^-1 mmol^-1] | ideal gas constant | 8.3145 [J K^-1 mol^-1] ideal gas constant | [125] |
| T | 298.150 [K] | temperature | 25 [°C] temperature | experimental condition |
| NA | 6.022e+23 [mol^-1] | avogadros number | 6.0221e23 [mol^-1] avogadros number | [125] |
| M_chl | 893.509 [g mol^-1] | molar mass of chlorophyll a | 893.509 [g mol^-1] molar mass of chlorophyll a (C55H72MgN4O5) | [126] |
| M_CO2 | 44.010 [g mol^-1] | molar mass of CO2 | 44.01 [g mol^-1] molar mass of CO2 (CO2) | [126] |
| DeltaG0_ATP | 30.600 [kJ mol^-1] | energy of ATP formation | 30.6 [kJ mol^-1] energy of ATP formation | [48] |
| | | Concentrations | | |
| PSIItot | 0.830 [mmol mol(Chl)^-1] | total concentration of photosystem II complexes | measured photosystems fluorescence at 77K | [56] |
| PSItot | 3.270 [mmol mol(Chl)^-1] | total concentration of photosystem I complexes | measured photosystems fluorescence at 77K | [56] |
| Q_tot | 13.000 [mmol mol(Chl)^-1] | total PHOTOACTIVE PQ concentration | 13 [1000(Chl)^-1] total PHOTOACTIVE PQ concentration (Khorobrykh2020) | [54] |
| PC_tot | 1.571 [mmol mol(Chl)^-1] | total concentration of plastocyanin (PC_ox + PC_red) | 22000 [cell^-1] total concentration of plastocyanin (PC_ox + PC_red) | [33] |
| Fd_tot | 3.597 [mmol mol(Chl)^-1] | total concentration of ferredoxin (Fd_ox + Fd_red) | 1.1 [unitless] ratio of total ferredoxin (Fd_ox + Fd_red) to PSI | [121] |
| NADP_tot | 26.805 [mmol mol(Chl)^-1] | total concentration of NADP species (NADP + NADPH) | 30 [nmol mg(Chl)^-1] total concentration of NADP species (NADP + NADPH) | [127] |
| NAD_tot | 11.169 [mmol mol(Chl)^-1] | total concentration of NAD species (NAD + NADH) | 50 [nM OD^-1] total concentration of NAD species (NAD + NADH) | [110] |
| AP_tot | 430.143 [mmol mol(Chl)^-1] | total concentration of adenosine species (ADP + ATP) | 400 [pmol (10^8 cells)^-1] cellular content of ATP | [111] |
| Pi_mol | 1.000e-02 [mmol mol(Chl)^-1] | molar concentration of phosphate | 1.000e-02 [mmol mol(Chl)^-1] molar concentration of phosphate | [48] |
| S | 35.000 [unitless] | salinity within a cell | 35 [unitless] salinity of sea water since no cell estimations could be found | [119] |
| cuvette_Chlconc | 0 [mmol(Chl) m^-3] | chlorophyll concentration in the measured sample, used for calculation of light attenuation | no light attenuation assumed if it wasn't measured in a particular experiment | estimated |
| | | Standard electrode potentials | | |
| E0_QA | -0.140 [V] | standard electrode potential of the reduction of PS2 plastoquinone A | -0.14 [V] midpoint potential of the reduction of PS2 plastoquinone A | [12] |
| E0_PQ | 0.533 [V] | standard electrode potential of the reduction of free plastoquinone | 0.12 [V] midpoint potential of the reduction of free plastoquinone | [12] |
| E0_PC | 0.350 [V] | standard electrode potential of the reduction of free plastocyanin | 0.35 [V] midpoint potential of the reduction of free plastocyanin | [12] |
| E0_P700 | 0.410 [V] | standard electrode potential of the reduction of the oxidized PS1 reaction center | 0.48 [V] midpoint potential of the reduction of the oxidized PS1 reaction center | [12] |
| E0_FA | -0.580 [V] | standard electrode potential of the reduction of PS1 iron-sulfur cluster A | -0.58 [V] midpoint potential of the reduction of PS1 iron-sulfur cluster A | [12] |
| E0_Fd | -0.410 [V] | standard electrode potential of the reduction of free ferredoxin | -0.41 [V] midpoint potential of the reduction of free ferredoxin | [12] |
| E0_NADP | -0.113 [V] | standard electrode potential of the reduction of NADP to NADPH | -0.32 [V] midpoint potential of the reduction of NADP to NADPH | [128] |
| E0_succinate/fumarate | 0.443 [V] | standard electrode potential of the reduction of fumarate to succinate | 0.03 [V] midpoint potential of the reduction of fumarate to succinate | [128] |

Table S3 **- continued**

| Parameter | Value | Description | Source value | Source |
|---|---|---|---|---|
| | | General reaction parameters | | |
| kH0 | 5.000e+08 [s^-1] | rate constant of (unregulated) excitation quenching by heat | 5e8 [s^-1] rate of (unregulated) excitation quenching by heat | [47] |
| kHst | 1.000e+09 [s^-1] | rate constant of state transition regulated excitation quenching by heat | 1e9 [s^-1] rate of qE regulated excitation quenching by heat | [47] |
| kF | 6.250e+08 [s^-1] | rate constant of excitation quenching by fluorescence | 6.25e8 [s^-1] rate of excitation quenching by fluorescence | [47] |
| k2 | 2.500e+09 [s^-1] | rate constant of excitation quenching by photochemistry | 5e9 [s^-1] rate of excitation quenching by photochemistry, decreased by factor 0.5 in manual fit to achieve Fm/F0 ca. 1.6 | [47, 129] |
| k_F1 | 1.000 [s^-1] | rate constant of excitation quenching via fluorescence | manually fitted to achieve low intensity PSI fluorescence | manually fitted |
| kPQred | 250.000 [mol(Chl) mmol^-1 s^-1] | rate constant of PQ reduction via PS2 | 250 [mol(Chl) mmol^-1 s^-1] rate of PQ reduction via PS2 | [48] |
| kPCox | 2.500e+03 [mol(Chl) mmol^-1 s^-1] | rate constant of PC oxidation via PS1 | 2500 [mol(Chl) mmol^-1 s^-1] rate of PC oxidation via PS1 | [48] |
| kFdred | 2.500e+05 [mol(Chl) mmol^-1 s^-1] | rate constant of Fd reduction via PS1 | 2.5e5 [mol(Chl) mmol^-1 s^-1] rate of Fd reduction via PS1 | [48] |
| k_Q | 1.755e+03 [mol(Chl) mmol^-1 s^-1] | rate constant of PC reduction by the cytochrome b6f complex | 1/10 [ms] reduction of half the total PC at the end of a high light pulse, increased by factor 10 in manual fit for full PQ oxidation in blue light | [130], manually fited |
| k_NDH | 3.136 [mol(Chl) mmol^-1 s^-1] | rate constant of PQ reduction by NDH-2 | 20 [umol(electrons) mg(Chl)^-1 h^-1] rate of electron flux through NDH-2 | [53] |
| k_SDH | 0.430 [mol(Chl) mmol^-1 s^-1] | rate constant of PQ reduction by SDH | 70 [umol(electrons) mg(Chl)^-1 h^-1] rate of electron flux through SDH | [53] |
| k_O2 | 12.926 [mol(Chl)^2 mmol^-2 s^-1] | rate constant of Fd oxidation by Flv 1/3 | 20 [umol(O2) mg(Chl)^-1 h^-1] rate of oxygen consumption by Flv 1/3 in dark-light transition | [131] |
| k_aa | 1.156 [mol(Chl)^2 mmol^-2 s^-1] | rate constant of oxygen reduction via COX | 4 [umol(O2) mg(Chl)^-1 h^-1] rate of oxygen reduction via COX, increased by factor 2 in manual fit to achieve a dark PQ reduction of ca. 20 % | [131], manually fitted |
| k_FN_fwd | 3.730 [mol(Chl) mmol^-1 s^-1] | rate constant of NADP reduction by FNR | 1/8 [ms] 8 ms time for reduction of half the NADP reduced during a saturating flash | [127] |
| k_FN_rev | 53.364 [mol(Chl)^3 mmol^-3 s^-1] | rate constant of reverse flux through FNR in darkness | 8.3 [umol(O2) mg(Chl)^-1 h^-1] rate of oxygen reduction in the dark, should roughly match the rate of electron flux trough FNR | [127] |
| kRespiration | 3.506e-06 [mol(Chl)^3 mmol^-3 s^-1] | rate constant of 3PGA oxidation and fumarate reduction by glycolysis and the TCA cycle | 8.3 [umol(O2) mg(Chl)^-1 h^-1] rate of oxygen reduction in the dark, reduced by factor 20 in manual fit to avoid over reduciton in darkness | [131] |
| kO2out | 4.025e+03 [s^-1] | rate constant of oxygen diffusion out of the cell | 62.05 [mmol mol(Chl)^-1 s^-1] O2 efflux out of the cell assuming an O2 membrane gradient of 0.064 uM | [108] |
| kCCM | 4.025e+03 [s^-1] | rate constant of CO2 diffusion into the cell, assumed to be identical to kO2out | estimated from kO2out | estimated |
| k_pass | 1.000e-02 [mmol mol(Chl)^-1 s^-1] | rate constant of protons leaking across the thylakoid membrane per delta pH | manually fitted to provide a low capacity flow dissipating the pH gradient | manually fitted |
| kATPsynth | 10.000 [s^-1] | rate constant of ATP synthesis | manually fitted to supply the CBB | manually fitted |
| kATPconsumption | 0.300 [s^-1] | rate constant of ATP consumption by processes other than the CBB | manually fitted to provide a low capacity flow consuming ATP | manually fitted |
| kNADHconsumption | 10.000 [s^-1] | rate constant of NADH consumption by processes other than NDH | manually fitted to provide a reasonable NAD+/NADH ratio | manually fitted |
| k_ox1 | 0.211 [mol(Chl)^2 mmol^-2 s^-1] | rate constant of oxygen reduction via bd-type (Cyd) terminal oxidases | 3 [umol(O2) mg(Chl)^-1 h^-1] approximate rate of oxygen reduction via bd-type (Cyd) terminal oxidases | [131] |
| KHillFdred | 39.062 [mmol^nHillFdred mol(Chl)^-nHillFdred] | Flv binding constant of Fd_red in Hill kinetics, assuming half activity around 2.5 mmol mol(Chl)^-1 | manually fitted to induce fast Flv activation at ca. 50 µmols m-2 s-1 white irradiance (Schuurmans, 2015) | manually fitted |
| nHillFdred | 4.000 [unitless] | Hill constant of Fd_red binding to Flv, assuming strong cooperative binding (see Brown2019) | manually fitted to induce fast Flv activation at ca. 50 µmols m-2 s-1 white irradiance ([25]) | manually fitted |
| vNQ_max | 50.000 [mmol mol(Chl)^-1 s^-1] | maximal rate of NDH-1 | manually fitted to achieve 30 % CET in Fig. 1b | manually fitted to Fig. 1b |
| KMNQ_Qox | 1.300 [mmol mol(Chl)^-1] | Michaelis constant for Q_ox reduction by NDH-1 | manually fitted to provide mostly constant NDH-1 flux into high light ([65]) | manually fitted |
| KMNQ_Fdred | 1.439 [mmol mol(Chl)^-1] | Michaelis constant for Fd_red oxidation by NDH-1 | manually fitted to provide mostly constant NDH-1 flux into high light ([65]) | manually fitted |
| | | Regulation parameters | | |
| kQuench | 2.000e-03 [mmol^-1 mol(Chl)^-1 s^-1] | rate constant of PS2 quenching by Q_red | manually fitted to Fig. 2a | manually fitted to Fig. 2a |
| kUnquench | 0.100 [s^-1] | rate constant of internal PS2 unquenching | manually fitted to Fig. 2a | manually fitted to Fig. 2a |
| KMUnquench | 0.200 [mmol^ mol(Chl)^-1] | binding constant of PS2 unquenching inhibition by Q_red | manually fitted to Fig. 2a | manually fitted to Fig. 2a |
| kCBBactivation | 3.466e-02 [s^-1] | rate constant of CBB activation by reduced ferredoxin | manually fitted to reproduce CBB activation within 2 mins in dark to light transition [55] | manually fitted |
| KMFdred | 0.300 [mmol mol(Chl)^-1] | Michaelis constant of CBB activation by reduced ferredoxin | manually fitted to achieve CBB activation under at ca. 10 µmol m-2 s-1 light where respiration becomes repressed [25] | manually fitted |
| OCPmax | 0.280 [unitless] | maximal fraction of PBS quenched by OCP | ca. 29% of PBS were quenched in WT with 220 µE m-2 s-1 blue green illumination, maximum assumed to be 10% higher, manually fitted to Fig. 2a | [132], manually fitted |
| kOCPactivation | 9.600e-05 [s^-1 (umol(Photons) m^-2 s^-1)^-1] | rate constant of OCP activation by absorbed light | manually fitted to Fig. 2a | manually fitted |
| kOCPdeactivation | 1.350e-03 [s^-1] | rate constant of constitutive OCP relaxation by thermal processes and FRP | manually fitted to Fig. 2a | manually fitted |
| | | CBB and PR parameters | | |
| vCBB_max | 51.301 [mmol mol(Chl)^-1 s^-1] | approximate maximal rate of the Calvin Benson Bassham cycle | 0.2e-15 [g(CO2) cell^-1 s^-1] approximate maximal CO2 uptake rate | [133] |
| vOxy_max | 16.059 [mmol mol(Chl)^-1 s^-1] | approximate RuBisCO oxygenation rate | 1 [s^-1] approximate RuBisCO oxygenation rate | [134] |
| KMATP | 24.088 [mmol mol(Chl)^-1] | Michaelis constant for ATP consumption in the CBB cycle | 0.1 [mmol l^-1] order of magnitude of KM_ATP for phosphoribulokinase (Wadano1998) and phosphoglyceratekinase | [135, 114] |
| KMNADPH | 18.066 [mmol mol(Chl)^-1] | Michaelis constant for NADPH consumption in the CBB cycle | 0.075 [mmol l^-1] approximate KM_NADPH for GAP2 in Synechocystis | [136] |
| KMCO2 | 72.264 [mmol mol(Chl)^-1] | Michaelis constant for CO2 consumption by cyanobacterial RuBisCO | 0.3 [mmol l^-1] order of magnitude for KM_CO2 of cyanobacterial RuBisCO | [134] |
| KICO2 | 72.264 [mmol l^-1] | inhibition constant of CO2 for O2 consumption by cyanobacterial RuBisCO, assumed equal to KMCO2 | 0.3 [mmol l^-1] order of magnitude for KM_CO2 of cyanobacterial RuBisCO | estimated |
| KMO2 | 240.880 [mmol mol(Chl)^-1] | Michaelis constant for O2 consumption by cyanobacterial RuBisCO | 1 [mmol l^-1] order of magnitude for KM_O2 of cyanobacterial RuBisCO | [134] |
| KIO2 | 240.880 [mmol mol(Chl)^-1] | inhibition constant of O2 for CO2 consumption by cyanobacterial RuBisCO, assumed equal to KMO2 | 1 [mmol l^-1] order of magnitude for KM_O2 of cyanobacterial RuBisCO | estimated |
| KMPGA | 0.100 [mmol mol(Chl)^-1] | Michaelis constant limiting oxygenation reactions for low 3PGA | manually fitted to not limit respiration or photorespiratioo | manually fitted |
| kPR | 1.523e-06 [mol(Chl)^4 mmol^-4 s^-1] | rate constant of (2-phospho)glycolate recycling into 3PGA | 4.80e-6 [umol min^-1 ug(Chl)^-1] estimated rate of photorespiration (glycolate appearance) under Air-like CO2 concentration | [112] |

Table S4: **Parameters used for light-adapted strains.** The parameters were experimentally measured by Zavřel *et al.* (2024) [56] or Rodrigues *et al.* (2023) [29] or inferred from such measurements. COlumns designate cells grown at a certain monochromatic light wavelength and intensity. The entries `chla`, `beta_carotene`, `allophycocyanin`, `phycocyanin` are part of the parameter `pigment_content`. `cuvette_Chlconc` was used for determining light attenuation. Default values are used in the case of empty cells. For parameter descriptions, see Table S3.

| | Zavrel (2024) | | | | | | | | Rodrigues (2023) | | | | | | | Unit |
|---|---|---|---|---|---|---|---|---|---|---|---|---|---|---|---|---|
| Wavelength<br>Light intensity | 435 nm<br>25 | 465 nm<br>25 | 495 nm<br>25 | 520 nm<br>25 | 555 nm<br>25 | 633 nm<br>25 | 663 nm<br>25 | 687 nm<br>25 | 405 nm<br>50 | 405 nm<br>100 | 450 nm<br>50 | 540 nm<br>50 | 540 nm<br>100 | 630 nm<br>50 | 630 nm<br>100 | µmol(photons) $m^{-2}\,s^{-1}$ |
| PBS_PS2 | 0.597 | 0.5639 | 0.5992 | 0.568 | 0.53 | 0.5117 | 0.5167 | 0.615 | | | | | | | | unitless |
| PBS_PS1 | 0.3049 | 0.2909 | 0.2921 | 0.3071 | 0.3552 | 0.3891 | 0.4034 | 0.2773 | | | | | | | | unitless |
| PBS_free | 0.09814 | 0.1452 | 0.1088 | 0.1249 | 0.1147 | 0.09921 | 0.07988 | 0.1076 | | | | | | | | unitless |
| PSIItot | 1.818 | 1.221 | 1.221 | 1.093 | 0.8449 | 0.8314 | 0.936 | 2.031 | | | | | | | | mmol mol(Chl)$^{-1}$ |
| PSItot | 3.03 | 3.175 | 3.175 | 3.207 | 3.267 | 3.27 | 3.245 | 2.979 | | | | | | | | mmol mol(Chl)$^{-1}$ |
| chla | 1 | 1 | 1 | 1 | 1 | 1 | 1 | 1 | 1 | 1 | 1 | 1 | 1 | 1 | 1 | mg(Pigment) mg(Chla)$^{-1}$ |
| beta_carotene | 0.1812 | 0.17 | 0.1723 | 0.1541 | 0.1418 | 0.1765 | 0.1729 | 0.1882 | 0.1437 | 0.1471 | 0.142 | 0.143 | 0.1412 | 0.1441 | 0.1442 | mg(Pigment) mg(Chla)$^{-1}$ |
| allophycocyanin | 0.3906 | 0.025 | 0.1064 | 0.2838 | 0.4937 | 1.118 | 0.9792 | 0.7255 | 1.2295 | 1.6152 | 0.7983 | 0.8497 | 0.7942 | 0.8154 | 1.097 | mg(Pigment) mg(Chla)$^{-1}$ |
| phycocyanin | 3.453 | 4.775 | 4.085 | 3.081 | 3.582 | 6.765 | 5.542 | 4.608 | 7.5436 | 9.449 | 5.3145 | 5.765 | 5.6851 | 5.4425 | 6.73 | mg(Pigment) mg(Chla)$^{-1}$ |
| cuvette_Chlconc | 0.6797 | 0.09158 | 0.2174 | 0.9016 | 1.452 | 0.9766 | 0.9554 | 0.9257 | | | | | | | | mmol(Chl) $m^{-1}$ |

The parameter $fCin$, representing the intracellular to extracellular partial pressure ratio as increased by the carbon concentrating mechanism, was varied in the simulation between 100 and 1000 and has been set to 1000 based on the fit to the measured by Benschop *et al.* [120] $CO_2$ fixation rate (Fig. S2).

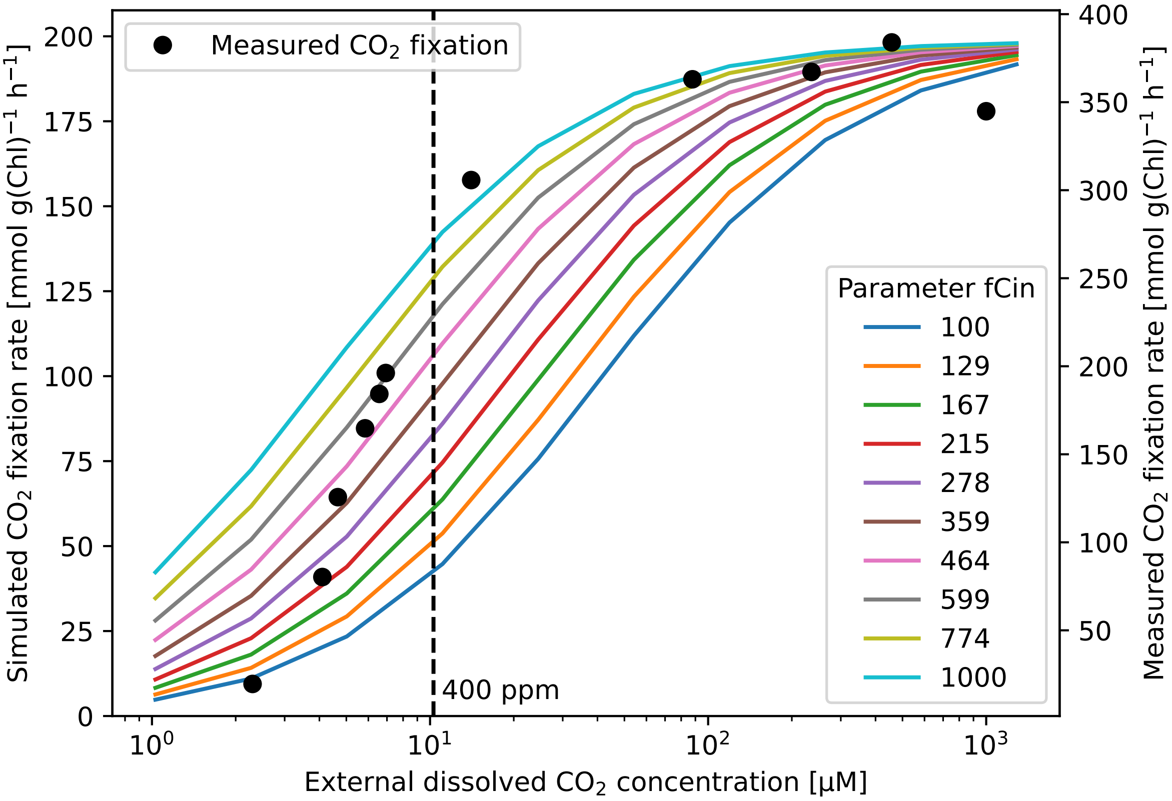

Fig. S2: **$O_2$ production under light intensity variation *in vivo* and *in vitro*.** The parameter $fCin$, representing the intracellular to extracellular partial pressure ratio as increased by the carbon concentrating mechanism, was varied in the simulation between 100 and 1000. Benschop *et al.* [120] measured oxygen evolution with 800 µmol(photons) $m^{-2}\,s^{-1}$ light of *Synechocystis* sp. PCC 6803 grown under 20 ppm $CO_2$ and varied the dissolved $CO_2$ in the medium ($C_i$). The simulation used a cool white LED. Our simulated $O_2$ evolution rates are ca. half of the measured rates. However, the rate dynamics in ambient air (400 ppm) or above are well reproduced with $fCin = 1000$. The model overestimates oxygen evolution at very low $C_i$ concentrations.

#### S2.2 Calculating model parameters from experimental data

In this work, we used measured pigment concentrations of the cyanobacterial samples as our input and we fit seven out of 23 parameters: Activation and inactivation rates of state transitions and OCP, the $K_m$ of state transition activation by reduced PQ, and the fluorescence contribution of PBS, to PAM-SP measurements of our previous publication [56]. We estimated the photosystem concentrations and attachment of PBS from 77K fluorescence excitation-emission maps [56] (see Table S4).

The 77K fluorescence ratio of PSI to PSII ($f_{I:II}^{Fluo}$) was found in the range of two to six, which is close to the physiological ratio [137]. Therefore we make the simplifying assumption that the fluorescence ratio is equal to the ratio of photosystem monomers. We first calculate the ratio of PSI to total photosystems ($f_I^{PS}$):

$$f_I^{PS} = \frac{1}{1 + {}^{1}/_{f_{I:II}^{Fluo}}} \tag{S62}$$

We use this fraction to estimate how much cellular chlorophyll is bound in PSI ($f_I^{Chl}$) or PSII ($f_{II}^{Chl}$), assuming all chlorophyll is present in photosystems

$$f_I^{Chl} = 1 - f_{II}^{Chl} = \frac{f_I^{PS} \cdot n_I^{Chl}}{f_I^{PS} \cdot n_I^{Chl} + f_{II}^{PS} \cdot n_{II}^{Chl}} \tag{S63}$$

with the number of chlorophyll per PSI ($n_I^{Chl} = 96$) and PSII ($n_{II}^{Chl} = 35$). The inverse of the photosystems' chlorophyll content gives us the theoretical maximum chlorophyll-specific concentration. Thus we can calculate the concentration of PSI trimers ($c_I$) and PSII dimers ($c_{II}$) as:

$$3c_I = c_{I,mono} = f_I^{Chl} \cdot \frac{1000}{n_I^{Chl}} \tag{S64}$$

$$2c_{II} = c_{II,mono} = f_{II}^{Chl} \cdot \frac{1000}{n_{II}^{Chl}} \tag{S65}$$

To determine PBS association, we calculated the approximate 77K fluorescence signal from PBS associated with either PSI, PSII or free PBS (for details see [56]). We assumed that the signal of free PBS was ten times higher than those bound to photosystems (compare [26]).

#### S2.3 Calculating the mean light in a light absorbing culture

For simulations of experimental data where the culture density was provided, we further calculate the light encountered by a mean cell ($I$) for each wavelength according to an integrated Lambert-Beer function [62] accounting for the decreasing irradiance at various depths ($L$) due to cellular absorption. While simulating PAM-SP experiments we have assumed $L = 1$cm, corresponding to the diameter of the used measuring cuvette. We further estimate the pigment concentrations in the cuvette $c_p$, The resulting vector $I$ has to take into account the culture's wavelength $\lambda$-specific absorption $a_{\lambda,tot}$ (normalized by volume), which we calculate from the absorption coefficients $a_{\lambda,p} \in A$ (normalized by chlorophyll, Equation 2):

$$a_{\lambda,tot} = \sum_p a_{\lambda,p} \cdot c_p \tag{S66}$$

$$i_\lambda = \frac{1}{L} \int_0^L i_{\lambda,0} \cdot e^{l \cdot a_{\lambda,tot}} dl \quad \text{with } i_\lambda \in I,\ i_{\lambda,0} \in I_0 \tag{S67}$$

If no light attenuation is assumed $I = I_0$.

Table S5: **State transition model parameters used in the model.** Each parameter is given with the lower and upper bound, where it was varied in the systematic perturbations.

| Parameter | Default value | Lower bound | Upper bound | Unit |
|---|---|---|---|---|
| kUnquench | 0.1 | 0.01 | 1 | mmol^-1 mol(Chl) s^-1 |
| kQuench | 2e-3 | 2e-2 | 2e-4 | mmol^-1 mol(Chl) s^-1 |
| KMUnquench | 0.2 | 0.01 | 0.3 | mmol mol(Chl)^-1 |
| kspill | 5e-3 | 5e-4 | 5e-2 | mmol^-1 mol(Chl) s^-1 |
| kunspill | 5e-4 | 5e-5 | 5e-3 | mmol^-1 mol(Chl) s^-1 |
| spillmax | 0.3 | 0.1 | 0.6 | unitless |
| kPBS_toPS1 | 5e-3 | 5e-4 | 5e-2 | mmol^-1 mol(Chl) s^-1 |
| kPBS_toPS2 | 1e-3 | 1e-4 | 1e-2 | mmol^-1 mol(Chl) s^-1 |
| PBS_PS1min | 0.25 | 0 | 0.5 | mmol mol(Chl)^-1 |
| PBS_PS2min | 0.35 | 0 | 0.5 | mmol mol(Chl)^-1 |
| kPBS_detach | 1e-4 | 1e-5 | 1e-3 | mmol^-1 mol(Chl) s^-1 |
| kPBS_attach | 1e-3 | 1e-4 | 1e-2 | mmol^-1 mol(Chl) s^-1 |
| PBS_freemax | 0.1 | 0.01 | 0.3 | unitless |

### S3 Additional model validation

#### S3.1 Dynamics of photoprotection

We have used our model to calculate fluorescence signal from cells grown under 633 633 nm. We have changed the input of the pigment composition and performed simulations using parameters fitted in Fig. 2a.

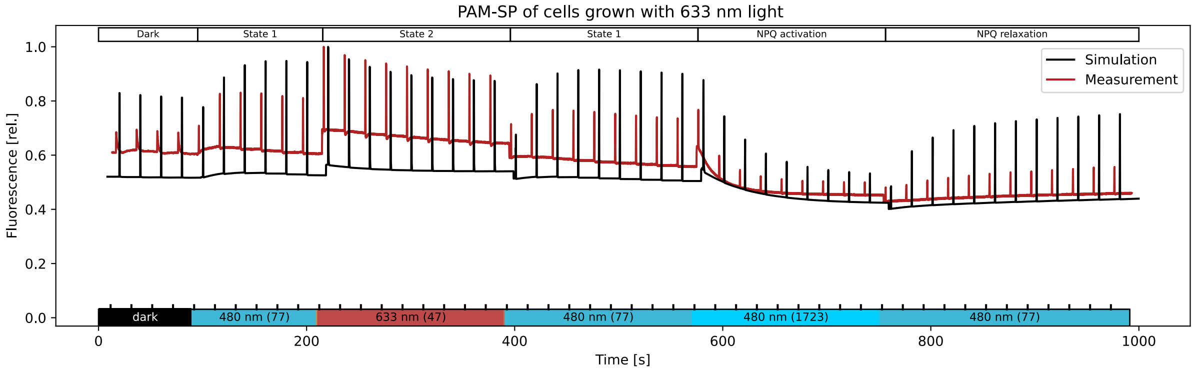

Fig. S3: Comparison of simulated (black) to measured (red) pulse amplitude modulation fluorescence dynamics during a saturation pulse light protocol. The simulation uses the parameters fitted in Fig. 2a. The experimental traces were measured in *Synechocystis* sp. PCC 6803 grown under 633 nm light (n=2) [56]. Simulations use the measured pigment contents and ambient $CO_2$ (400 ppm). The shown light protocol includes several different light wavelengths and intensities to trigger a response from respective photosynthetic electron transport chain components. By monitoring cell responses to these light conditions, we captured light responses via state transitions and non-photochemical quenching and relaxation (as described in the upper bar). We calculated light attenuation in the culture using Equation S67 with the measured pigment concentrations and sample chlorophyll content in Table S4.

#### S3.2 Oxygen production rate

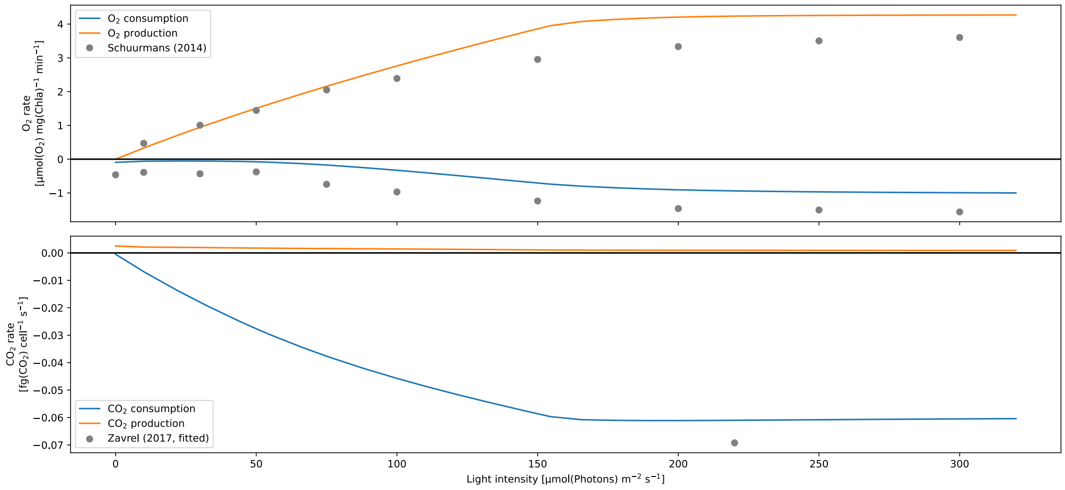

Fig. S4: **Rates of change in $O_2$ and $CO_2$.** Our model reproduces oxygen production and consumption (respiration + three terminal oxidases) rates for increasing light intensities, as compared to measured rates digitized from graphs by Schuurmans *et al.* (2014) [64]. Data points are from measurements of cells with 625 nm illumination with 50 mmol $L^{-1}$ $NaHCO_3$. Small variations in the data and the lack of error bars are due to the digitization process. The simulated carbon fixation rates are displayed with the measurement used for parameterization [133]. Simulations were run using a 625 nm light (gaussian LED, $\sigma = 10$ nm). We calculated light attenuation in the culture using Equation S67 with default pigment concentrations and a constant 2 mg $L^{-1}$ sample chlorophyll concentration.

#### S3.3 Internal PSII states

As discussed in the main text we calculated the fraction of open PSII for increasing light intensities and compared it to measurements [65]. Although our response curve is less sensitive to increasing light, and PSII are still open for light intensities higher than reported 300 µmol(photons) $m^{-2}$ $s^{-1}$ we observe the expected exponential reduction of the fraction of open PSII with increasing light intensity.

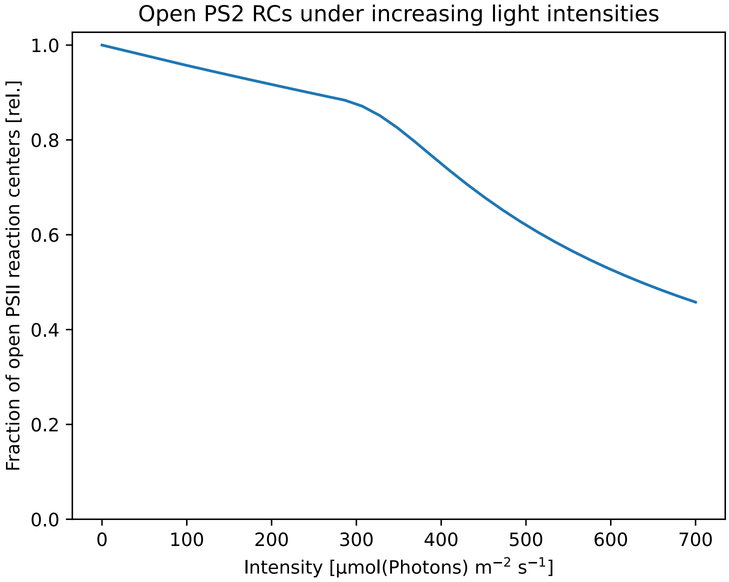

Fig. S5: **Steady-state fraction of open PSII reaction centers und different light intensities.** The model was simulated to steady-state under illumination with a fluorescence lamp spectrum at intensities between 0.1 and 700 µmol(photons) $m^{-2}$ $s^{-1}$. The open fraction was calculated as the fraction of PSII in non-reduced states $B_0$ and $B_1$ [47] (see Fig. S1).

#### S3.4 Activation of OCP in blue light

With our model analysis we could identify the activation of OCP under blue light saturation pulses (Fig. S6).

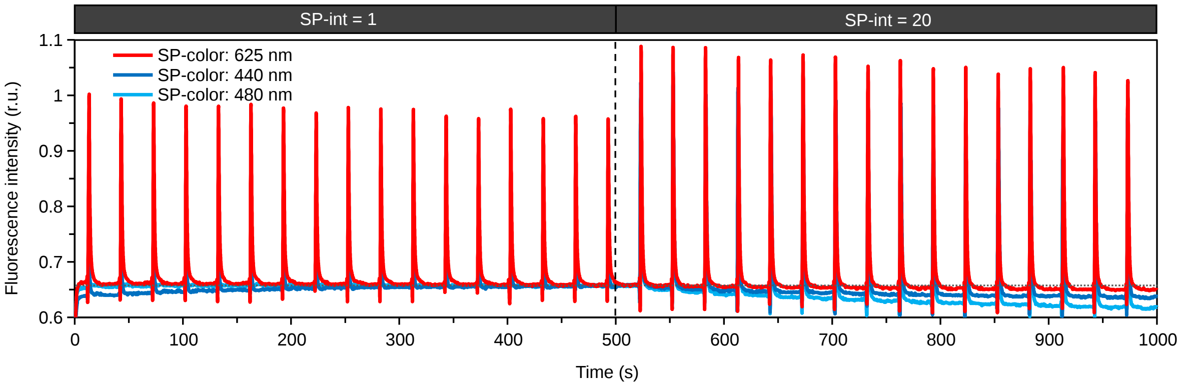

Fig. S6: **Light pulses of different wavelengths differ in triggering fluorescence quenching.** The measurements were performed with Multi-Color PAM (Walz, Effeltrich, Germany). Low-intensity pulses (SP-Int=1) affect the steady-state fluorescence (F) only weakly. With each pulse of 440 nm and 480 nm light, however, the F level decreases stepwise, pointing at fluorescence quenching, possibly through Orange Carotenoid Protein (OCP). The culture of *Synechocystis* sp. PCC 6803 was pre-cultivated in a conical flask on a shaker under cool white light (30 µmol(photons) $m^{-2}$ $s^{-1}$) at 23 °C to OD750=0.2 (measured with Shimadzu UV-Vis 2600 spectrophotometer, Shimadzu, Kyoto, Japan). For the measurement, 1.5 mL culture was transferred to a quartz cuvette and dark-acclimated for 5 min prior to each measurement. During the measurement, a custom-made protocol was used with the following settings: Analysis mode: SP analysis; AL off; SP-int=1 (500 s) / 20 (500 s); SP-color= 440 nm / 480 nm / 625 nm; ML-color=625 nm.

### S4 Additional analysis

#### S4.1 Mutant analysis

We extended our analysis of the impact of light intensity on the wild type and three mutants by additionally calculating their NPQ and photosynthesis yield (Fig. S7) and fraction of reduced pools, lumenal and stromal pH, and main fluxes (Fig. S8).

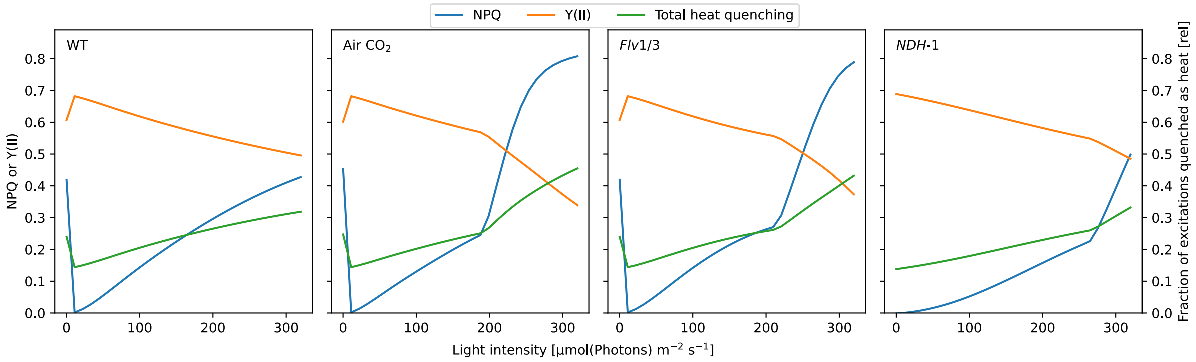

Fig. S7: **Calculated NPQ and photosynthetic yield (Y(II)) and the fraction of total excitations quenched as heat for four *in silico* lines** representing the wild type (WT) in saturating $CO_2$ and ambient air $CO_2$ (400 ppm), a flavodiiron (*Flv*1/3) and NAD(P)H Dehydrogenase-like complex 1 (NDH-1) knockout mutant. The electron fluxes were calculated according to subsection S1.7. The models were simulated to steady state for light intensities between 0.1 µmol(photons) $m^{-2}\,s^{-1}$ and 300 µmol(photons) $m^{-2}\,s^{-1}$. Modeled conditions as in Fig. 1.

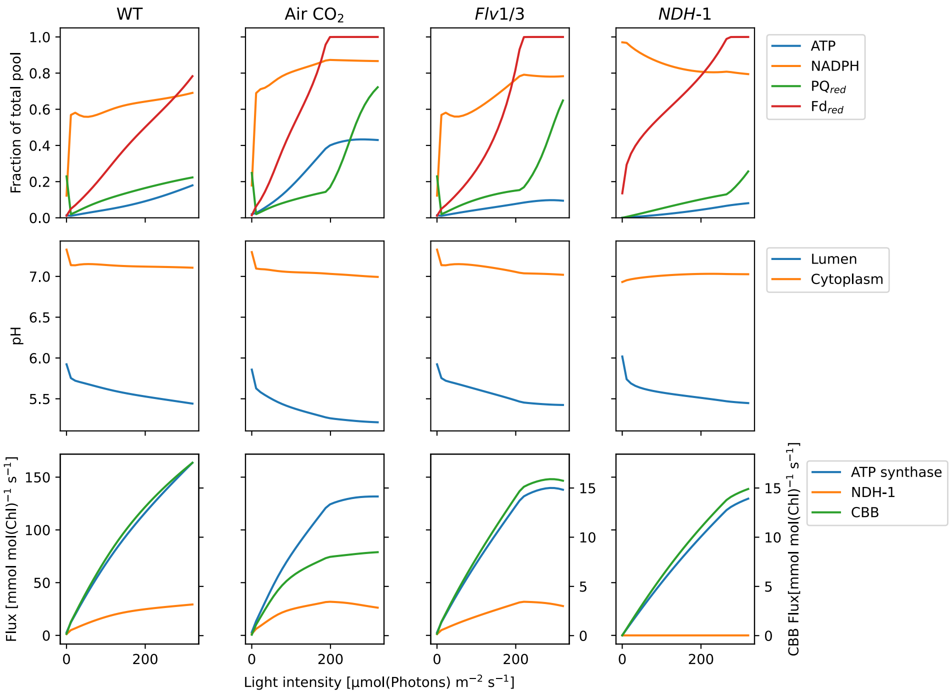

Fig. S8: **Simulated fraction of reduced pools, lumenal and stromal pH and fluxes through ATP synthase, carbon fixation (CBB) and cyclic electron flow (NDH) for four *in silico* lines** representing the wild type (WT) in saturating $CO_2$ and ambient air $CO_2$ level (400 ppm), a flavodiiron (*Flv*1/3) and NAD(P)H Dehydrogenase-like complex 1 (NDH-1) knockout mutant. The levels and production fluxes of ATP and NADPH are shown as the primary output metabolites of photosynthesis, next to central redox carriers. For better visibility, the CBB flux is rescaled to the rightward axis in the bottom plots.

#### S4.2 Metabolic Control Analysis (MCA)

Using metabolic control analysis (MCA), we quantified the control distribution in the system for different light intensities. We divided the set of all reactions into four non-overlapping sets belonging to: Light-driven electron flow: PSII, $Cb_6f$, PSI, NDH-1, ATPS, FNR; Respiration: SDH, NDH-2, "respiration" (sugar catabolism); RuBisCO reactions: CBB, Oxy; Terminal oxidases: Flv, Cyd (bb-type), COX ($aa_3$-type).

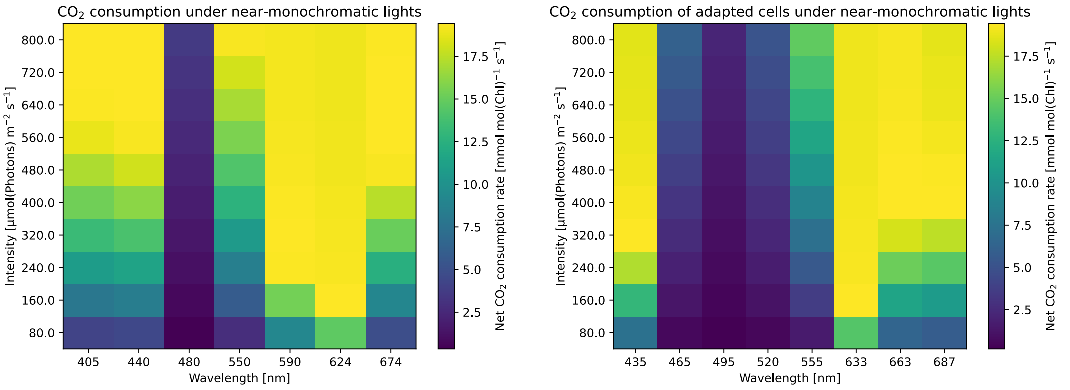

Fig. S9: **Simulated steady-state rates of $CO_2$ fixation under monochromatic lights with varying intensites.** Left) Simulations with default pigment composition. Right) Simulations with pigment compositions of *Synechocystis* sp. PCC 6803 grown under the respective light color. The adapted models were parameterized using pigment, photosystems, and PBS measurements. The models were then simulated to steady-state with the respective light condition, and the $CO_2$ consumption is shown. The $CO_2$ fixation rate is the lowest between 465 and 555 nm compared to all other tested conditions. Under 633 nm light, the highest $CO_2$ fixation rate is reached at the lowest intensity. Compared to the unadapted simulations, the efficient usage of red light and inefficient usage of blue light is more pronounced.

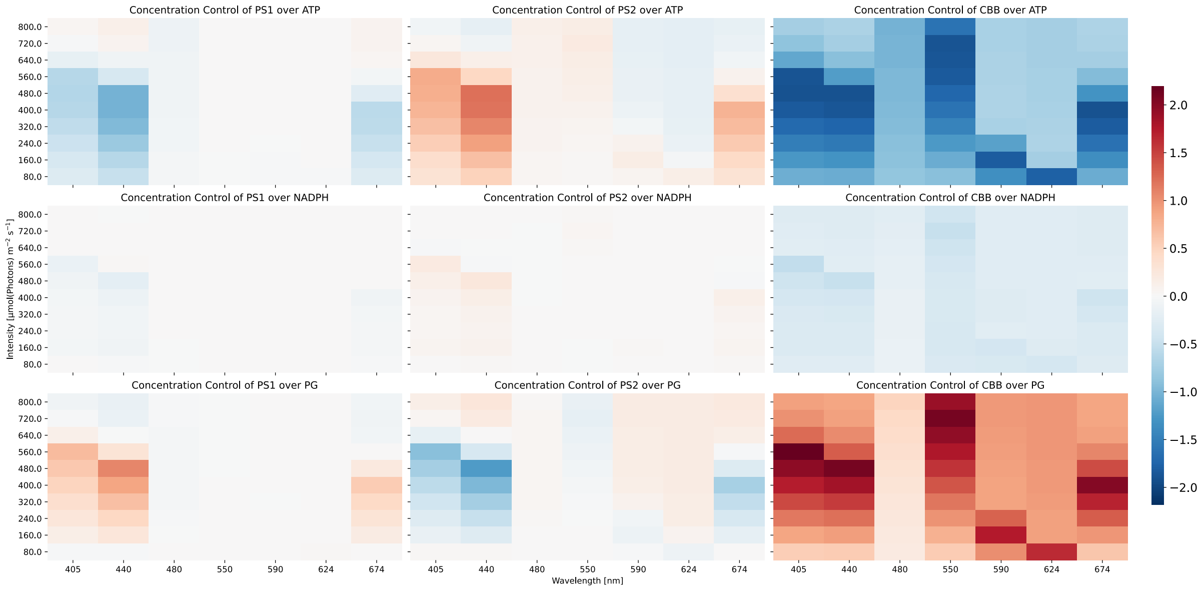

Fig. S10: **Control of photosystems and the CBB on the concentrations of ATP, NADPH, and 2PG.** We simulated the model under the lights in Fig. 5a with a range of intensities from 80 to 800 µmol(photons) $m^{-2}\,s^{-1}$ to steady-state. By varying the photosystem concentrations and the maximal rate of the CBB by ± 1 %, we quantified their control on the metabolite concentrations. More positive/negative values signify a stronger positive/negative control of the respective ETC component. The photosystems control is generally highest under illumination within the chlorophyll absorption spectrum (405 nm, 440 nm and 674 nm), with PSII also having control in the red spectrum. The CBB has a generally high control until a critical light intensity is reached. Increasing PSI or CBB flux generally lowers ATP and NADPH concentrations and increased 2PG, PSII has the opposite effect. At high light, some control relationships become inverted.

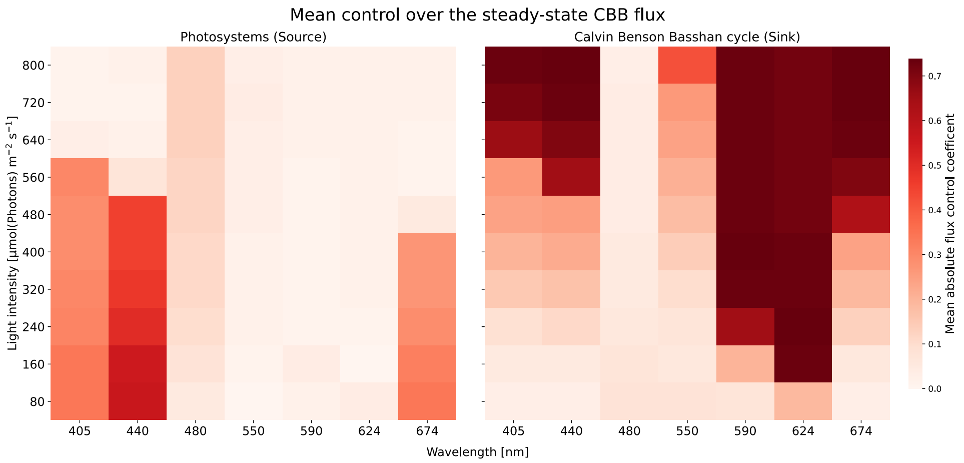

Fig. S11: **Results of metabolic control analysis under illumination with near-monochromatic Gaussian LEDs.** We simulated the model under the lights in Fig. 5a with a range of intensities from 80 to 800 µmol(photons) $m^{-2}\,s^{-1}$ to steady-state. By varying the photosystem concentrations and the maximal rate of the CBB by ± 1 %, we quantified their control on the CBB flux. Plots show the absolute control coefficients. The left graph shows the mean of both photosystems. Higher values signify stronger pathway control.

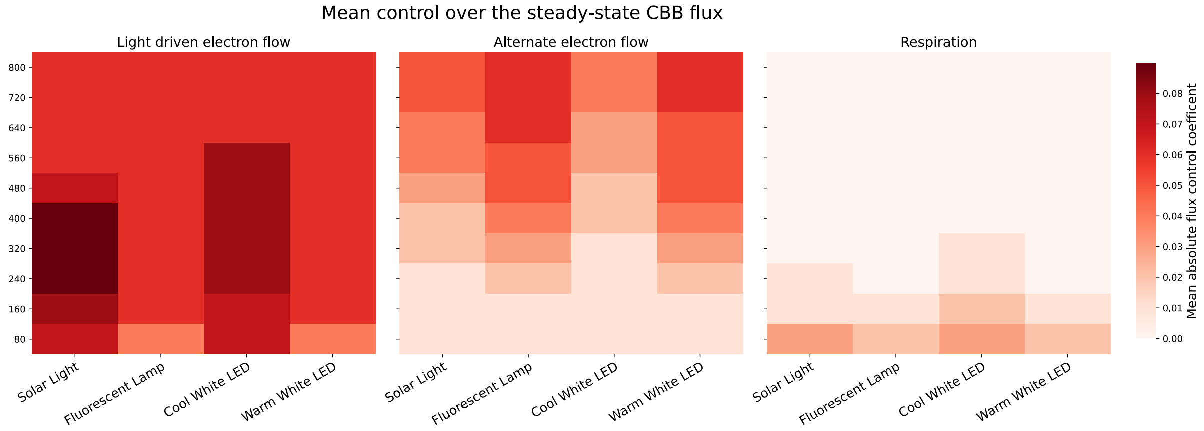

Fig. S12: **Results of Metabolic Control Analysis (MCA) performed for given light sources** in Fig. 3a with a range of intensities from 80 to 800 µmol(photons) $m^{-2}\,s^{-1}$ to steady-state. By varying the protein concentration, maximal velocity, or rate constant of a reaction by ± 1 %, we quantified their control on the CBB flux by calculating flux control coefficients. We display the absolutes of control coefficients as means within the following electron pathways: light-driven (PSI, PSII, Cytochrome $b_6f$ complex, NDH-1, FNR), alternate (Flv, Cytochrome bd quinol oxidase (Cyd), Cytochrome c oxidase), and respiration (lumped respiration, Succinate Dehydrogenase, NDH-2). Higher values signify stronger pathway control.

#### S4.3 Isoprene production

We were provided with the experimental data from [29] and we adapted our model to their measured pigment concentrations by changing three parameters, and found that the simulated isoprene productions follow their measured growth rate Fig. S13.

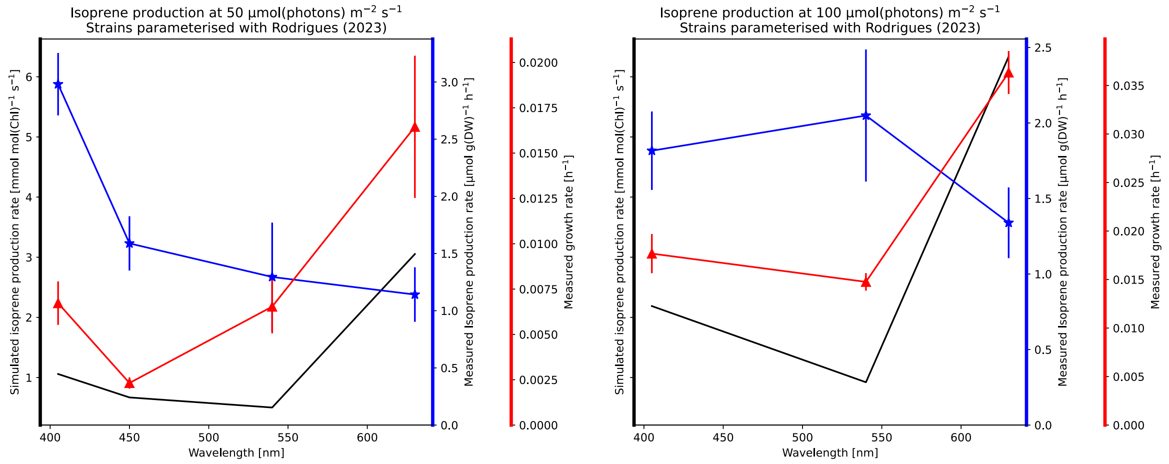

Fig. S13: **Comparison of measured isoprene production and simulated production capacity under light variation**. We show the growth and isoprene production rates measured under monochromatic illuminations (405 nm, 450 nm, 540 nm and 630 nm at 50 or 100 µmol(photons) $m^{-2}\,s^{-1}$) by Rodrigues *et al.* [29]. The model was adapted to the measured pigment composition (see Table S4) and simulated to steady-state under the respective monochromatic lights. We added a sink reaction to the model, consuming energy carriers in the ratio corresponding to isoprene (19 ATP, 11 NADPH, 4 $Fd_{red}$), including the cost of carbon fixation. We disabled the CBB reaction and limited ATP and NADPH concentrations to 95 % of their total pools. Simulations used a Gaussian LED ($\sigma$ =10 nm).
